# MglA functions as a three-state GTPase to control movement reversals of *Myxococcus xanthus*

**DOI:** 10.1101/577387

**Authors:** Christian Galicia, Sébastien Lhospice, Paloma Fernández Varela, Stefano Trapani, Wenhua Zhang, Jorge Navaza, Tâm Mignot, Jacqueline Cherfils

## Abstract

In *Myxococcus xanthus*, directed movement is controlled by inter-dependent pole-to-pole oscillations of the small GTPase MglA, its GAP MglB and the RomR protein. However, these proteins have strikingly different oscillatory regimes such that MglA is segregated from MglB and RomR at reversal activation. The molecular mechanism whereby information is exchanged between the lagging and leading poles resulting in MglA detachment from the leading pole during reversals has remained unknown. Here, we show that MglA has two GTP-bound forms, one of which is insensitive to MglB (MglA-GTP*) and is re-sensitized to MglB by a feedback mechanism operated by MglA-GDP. By identifying the region of MglB that is critical for its association to the lagging pole, we demonstrate that MglA-GTP* is functional *in vivo*. These data suggest that MglA-GDP acts as a soluble messenger to convert polar MglA-GTP* into a diffusible MglA-GTP species, explaining MglA re-localization to the opposite pole during reversals.

## Introduction

Small GTPases function as molecular switches in all living organisms by alternating between an inactive GDP-bound state and a GTP-bound state that recruits effectors (reviewed in (Cherfils and Zeghouf, 2011)). In general, GDP/GTP alternation is finely controlled by guanine nucleotide exchange factors (GEFs), which stimulate GDP/GTP exchange, and by GTPase-activating proteins or GAPs, which stimulate GTP hydrolysis (reviewed in (Cherfils and Zeghouf, 2013)). Polarity of migration controlled by small GTPases in prominent in eukaryotes (reviewed in (Charest and Firtel, 2007)), and has also been discovered in *Myxococcus xanthus*, a gram-negative deltaproteobacteria that moves by gliding on solid surfaces under the control of the small GTPase MglA and its GAP MglB (reviewed in (Treuner-Lange and Sogaard-Andersen, 2014)). *M. xanthus* uses two distinct motility machineries which operate in distinct conditions. The Social-motility (S-motility) complex, formed by so-called Type-IV pili, allows the collective motion of large groups of cells (reviewed in (Mauriello and Zusman, 2007)). In contrast, single motile cells at the periphery of the colony are propelled by the Adventurous-motility (A-motility) system comprised of the Agl-Glt complex (reviewed in (Islam and Mignot, 2015)). Remarkably, both systems are assembled at the bacterial pole under the control of a single small GTPase, MglA (Zhang et al., 2010, Leonardy et al., 2010). How MglA controls activity of the S-motility system remains to be established. In the case of A-motility, MglA-GTP assembles the Agl-Glt complex and recruits its components directly to the MreB actin cytoskeleton, thus traveling towards the lagging pole where the system is disassembled by inactivation of MglA by MglB (Treuner-Lange et al., 2015, Faure et al., 2016). In motile *M. xanthus* cells, MglA-GTP accumulates at the leading pole where it activates motility, while MglA-GDP is found diffusely in the cytosol (Zhang et al., 2010, Leonardy et al., 2010, Miertzschke et al., 2011). This localization pattern arises due to GAP activity of MglB, which is located at the lagging pole where it inactivates MglA-GTP and depletes it from this pole (Zhang et al., 2010, Leonardy et al., 2010). The ability of MglA to hydrolyze GTP is essential for motility regulation. Notably, locking MglA in the GTP-bound form by mutation of residues that are needed for GTP hydrolysis (e.g. Arg 53 or Gln82) or deletion of MglB provokes a characteristic pendulum A-motility regime, in which cells move by exactly one cell length between reversals (Zhang et al., 2010, Treuner-Lange et al., 2015). This remarkable behavior is due to impaired Agl-Glt disassembly at the lagging pole, enabling the complex to continue movement in the opposite direction. Thus far, whether regeneration of MglA-GTP from MglA-GDP at the new leading pole is spontaneous or requires an unknown GEF to stimulate GDP/GTP exchange has remained unknown.

Inversions of the direction of movement (reversals) is the inter-dependent pole-to-pole oscillations of MglA, MglB and another protein, RomR protein, which are controlled by signals transduced via the Frz receptor-kinase complex (reviewed in (Mercier and Mignot, 2016)). During movement, MglA and MglB remain segregated at opposite poles, while RomR steadily relocates from the leading pole to the lagging pole, with the duration of its complete re-localization defining the minimal period between reversals (Guzzo et al., 2018). Once RomR has fully relocated to the lagging pole, signals depending on phosphorylated FrzX, a substrate of the FrzE kinase that functions as a gate at the lagging pole, provoke the rapid detachment of MglA and its re-localization to the opposite pole, resulting in its transient co-localization with MglB. This co-localization correlates with a pause in movement, during which MglB rapidly relocates to the opposite pole, eventually allowing movement to resume in the opposite direction. Importantly, the slow re-localization of RomR is not regulated by signals and starts as soon as the cell undergoes a reversal. This regulatory design functions like a so-called gated relaxation oscillator, allowing the motility switch to adjust to signal levels (Guzzo et al., 2018). Intriguingly, although MglA, MglB and RomR bind directly to each other and RomR has been identified as a localization factor for MglA (Keilberg et al., 2012, Zhang et al., 2012), MglA-GTP is entirely segregated from MglB and RomR after RomR has re-localized to the lagging pole. Therefore, the persistence of MglA at the leading pole requires an as yet elusive mechanism. Likewise, how information is exchanged between the lagging and leading poles such as to detach MglA at the onset of a reversal, allowing its transient colocalization with MglB and subsequent inactivation, is not understood.

MglA and MglB are conserved in a large number of bacteria, where they are often encoded by a single operon (Wuichet and Sogaard-Andersen, 2014). Structural analysis of MglA-GDP from the extremophile *Thermus thermophilus* and its complex with MglB highlighted intriguing features not seen in eukaryotic small GTPases (Miertzschke et al., 2011). Notably, the switch 1 region, one of the two regions that bind nucleotides, undergoes a twisted 3-residue register shift between the inactive and the GAP-bound states, which moves catalytic residues in and out of register to stimulate GTP hydrolysis. Likewise, the switch 2 region, which in eukaryotes undergoes disorder-to-order transitions upon binding of GTP (reviewed in (Wittinghofer and Vetter, 2011)), adopts a well-ordered conformation in MglA-GDP that occludes the binding site of the γ-phosphate of GTP and is displaced by 5 Å to bind GTP in the MglA-MglB complex (Miertzschke et al., 2011). The structure of unbound MglA-GTP is currently unknown, leaving open the question of whether the substantial remodeling at the switch 1 and switch 2 is solely due to GTP or is promoted by MglB. MglB also differs from eukaryotic GAPs, which insert residues, often an arginine, near GTP to complete the catalytic sphere and stabilize the transition state (reviewed in (Cherfils and Zeghouf, 2013)). Instead, MglB forms no direct contact with GTP but indirectly positions a switch 1-borne arginine in a catalytic position (Miertzschke et al., 2011), in a manner reminiscent of the RGS GAPs of eukaryotic heterotrimeric G proteins (Tesmer et al., 1997). While important features of the GDP/GTP cycle of MglA were established by these structures, their roles in maintaining MglA-GTP at the leading pole during movement and allowing its rapid detachment at the onset of a reversal, is currently not understood.

In this study, we uncovered properties of MglA that provide important insight into these issues, by combining the determination of the full structural cycle of *M. xanthus* MglA and MglB, GAP kinetics reconstituted from purified proteins and *in vivo* motility assays. Our study reveals that MglA has in fact three major functional states, one bound to GDP and two bound to GTP. One of the GTP-bound states (MglA-GTP* hereafter) has a mixed inactive/active conformation, which renders it insensitive to GTP hydrolysis by MglB. Remarkably, MglA-GTP* can be reverted to the MglB-sensitive MglA-GTP form by MglA-GDP, uncovering a feedback mechanism mediated by MglB. By designing a diffusible MglB mutant able to reach all MglA-GTP species in the bacteria, which failed to displace the polar MglA-GTP species, we demonstrate that the MglB-resistant MglA-GTP* species exists *in vivo*. Together, our findings suggest that the MglA-GTP cluster located at the leading pole is composed of two MglA-GTP populations, including a major MglA-GTP* species that is resistant to GTP hydrolysis and a minor diffusible MglA-GTP species that can interact with the motility machineries and be inactivated by MglB at the lagging pole. When cells reverse, a rapid increase of the MglA-GDP pool could therefore convert the entire polar pool of MglA-GTP* into a diffusible MglB-sensitive MglA-GTP form, and thus explaing how the decision to reverse is conveyed from the lagging pole to the leading pole.

## Results

### MglA has a three-state GDP/GTP structural switch

To gain insight into the structural landscape of *M. xanthus* MglA, we determined the crystal structures of MglA in different nucleotide states, which uncovered that MglA can adopt three conformations: inactive MglA-GDP, active MglA-GTPγS, and a novel conformation combining inactive and active features **(Table 1 and S1, Figures 1A to 1H).** In MglA-GDP, the inactive switch 1 is retracted, which positions the catalytic arginine finger (Arg 53) remote from the nucleotide, and the switch 2 has an autoinhibitory conformation that occludes the binding sites of GTP and Mg^2+^, as seen in *T. thermophilus* MglA-GDP (Miertzschke et al., 2011) (**Figures 1A and 1B**). Binding of GTPγS induces a twisted 3-residue shift of the switch 1 and remodels the switch 2 to create space for the γ-phosphate, similar to MglA in the *T. thermophilus* MglA-MglB complex (Miertzschke et al., 2011) (**Figures 1C and 1D**). The shift of the switch 1 results in the elongation of the arginine finger loop (residues 44-58) (**Figure 1E**), but Arg53 remains outside the nucleotide-binding site. Displacement of the switch 2 also allows Mg^2+^ to bind with an incomplete coordination that lacks the conserved Mg^2+^-binding threonine (Thr54) located in the arginine finger loop **(Figure 1D and 1F).** Remarkably, we trapped MglA in a mixed state in which the switch 1 is in the inactive conformation and the switch 2 is in the active conformation, a combination never observed in a small GTPase before **(Figures 1E, 1F, 1G and 1H).** The nucleotide-binding site contains GDP and a ligand tentatively ascribed to a phosphate/sulfate with two partially occupied positions. There is no Mg^2+^ present in this structure, whose binding is hindered by the retracted switch 1. Overlay with MglA-GTPγS shows that GTP can readily be accommodated in the mixed MglA conformation (**Figure 1I**). Together, our crystallographic analysis reveals that MglA is a three-state GTPase through its ability to adopt a mixed conformation that combines a retracted switch 1 and an active switch 2.

**Table 1.**
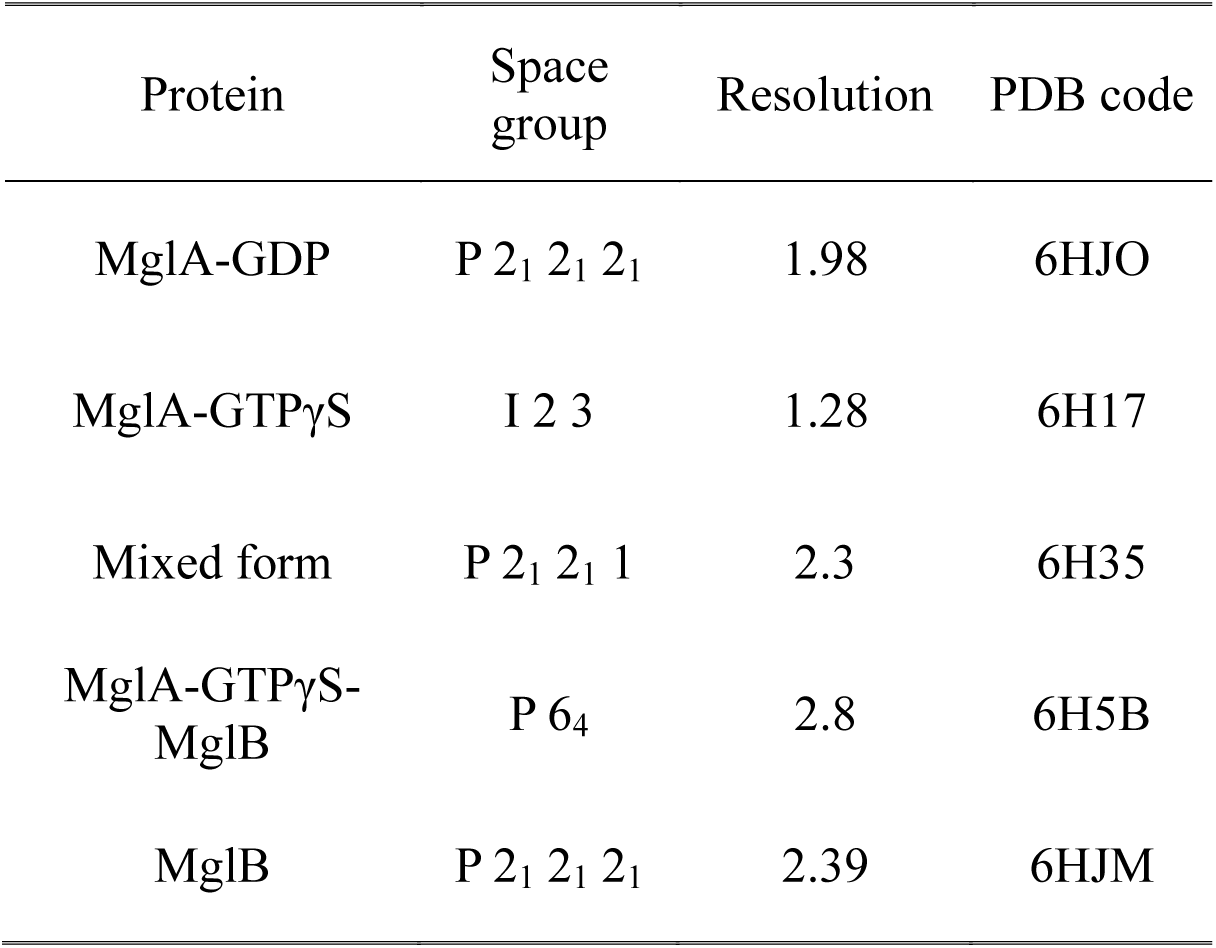
Crystal structures determined in this study

| Protein | Space group | Resolution | PDB code |
| --- | --- | --- | --- |
| MglA-GDP | P 2<sub>1</sub> 2<sub>1</sub> 2<sub>1</sub> | 1.98 | 6HJO |
| MglA-GTPγS | I 2 3 | 1.28 | 6H17 |
| Mixed form | P 2<sub>1</sub> 2<sub>1</sub> 1 | 2.3 | 6H35 |
| MglA-GTPγS-MglB | P 6<sub>4</sub> | 2.8 | 6H5B |
| MglB | P 2<sub>1</sub> 2<sub>1</sub> 2<sub>1</sub> | 2.39 | 6HJM |

**Figure 1.**
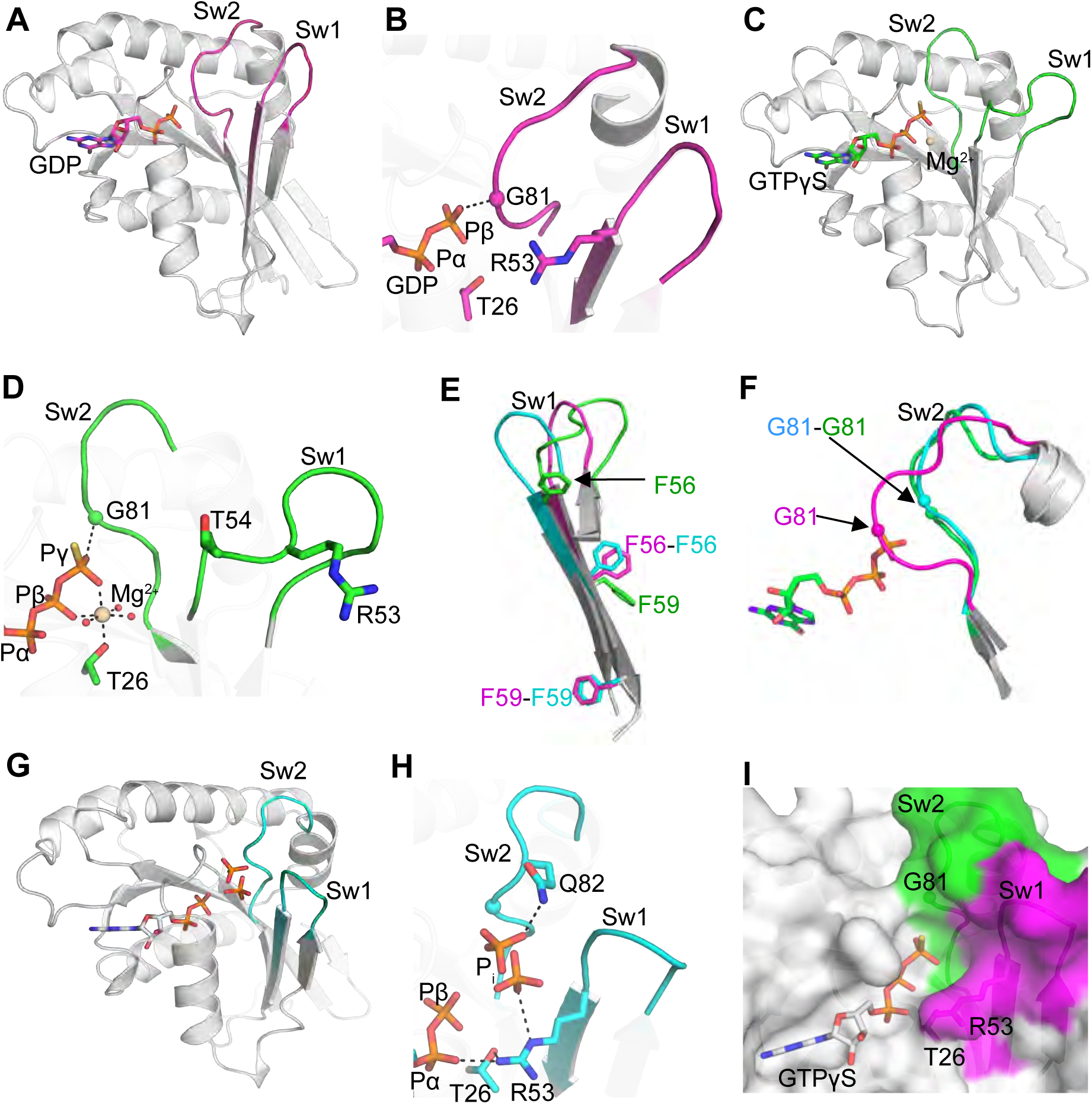
MglA has a 3-state GDP/GTP switch. **A-** Structure of MglA-GDP. The switch 1 is retracted and the switch 2 is autoinhibitory. **B-** Close-up view of the phosphate- and metal-binding sites of MglA-GDP. The arginine finger is remote from the nucleotide and Pro80 and Gly81 from the switch 2 occlude the Mg^2+^ and γ-phosphate binding sites. **C-** Structure of MglA-GTPγS-Mg^2+^. The switch 1 and the switch 2 are in active conformations. **D-** Close-up view of the phosphate- and metal-binding sites in MglA-GTPγS. The arginine finger is remote from the nucleotide and the coordination sphere of Mg^2+^ is incomplete. **E-** Overlay of the switch 1 from MglA-GDP, MglA-GTPγS and the mixed MglA form. The twisted 3-residue shift in the GTPγS-bound form is highlighted by the positions of F56 and F59. Note the flexibility of the switch 1 loop. **F-** Overlay of the switch 2 in MglA-GDP, MglA-GTPγS and the mixed MglA form, showing the large difference near Gly 81. Only GTPγS is shown for clarity. **G-** The mixed MglA form features a retracted switch 1 and an active switch 2 conformations. The structure contains GDP and phosphate or sulfate. **H-** Close-up view of the phosphate- and metal-binding sites of the mixed MglA form. The partially occupied sites of the phosphate/sulfate are shown. **I-** The mixed MglA form can accommodate GTP. Overlay of GTPγS taken from the MglA-GTPγS structure onto the mixed MglA structure. The surface of the nucleotide-binding site is shown as a surface, with the retracted switch 1 in pink and the active switch 2 in green. Color-coding for all panels except 1I is: GDP-bound; pink; GTPγS-bound: green; mixed form: blue. Hydrogen bonds are shown as dotted lines. Mg^2+^ is shown in beige. Sw1: switch 1. Sw2: switch 2.

### Crystal and solution structural analysis of MglB

To complete the MglA GDP/GTP structural cycle, we determined the crystal structures of its GAP MglB in unbound form and in complex with MglA-GTPγS **(Tables 1 and S1)**. The MglB crystal contains 20 molecules in the asymmetric unit, for which we applied an original strategy based on a low-resolution envelope derived from another weakly diffracting crystal form, making it one of the molecular replacement structure determination with the largest number of independent molecules to date (see Methods). The 20 monomers are arranged as 5 tetramers **(Figure S1A)**, which resemble tetramers seen in *T. thermophilus* MglB crystal structures in which the MglA-binding site is masked (Miertzschke et al., 2011). However, SEC-MALS analysis shows that the major species in solution is a dimer **(Figure S1B).** As previously observed for *T. thermophilus* MglB, *M. xanthus* MglB subunits have a roadblock fold (**Figure S1A**), which is frequently encountered in regulators of small GTPases (reviewed in (Cherfils, 2017)). The C-terminus (residues 133 to 165) is not visible in any of the 20 copies of the molecule, indicating that it is highly flexible. We used SEC-SAXS to analyze the structure of this segment in solution **(Figures 2A, S1C and S1D).** The dimensionless Kratky plot is representative of a globular protein with a significant proportion of flexible regions **(Figure S1E).** The maximal dimension (D_max_) of 96 Å is significantly larger than the D_max_ of the MglB core that is visible in the crystal (73Å), with the SAXS envelope showing additional volumes extending the dimeric core on each side **(Figure 2B).** Combining the SAXS and crystallographic data, the two C-terminii missing from the crystal structures could be modeled as intrinsically disordered peptides, yielding an excellent fit with the experimental data (χ^2^=1.22) **(Figure 2C and S1F).** Thus, the C-terminus is a flexible segment that enlarges the volume occupied by MglB. Its sequence is conserved across species (**Figure S1J**), which suggests that it plays a role in MglB functions, possibly exploiting its flexibility.

**Figure 2.**
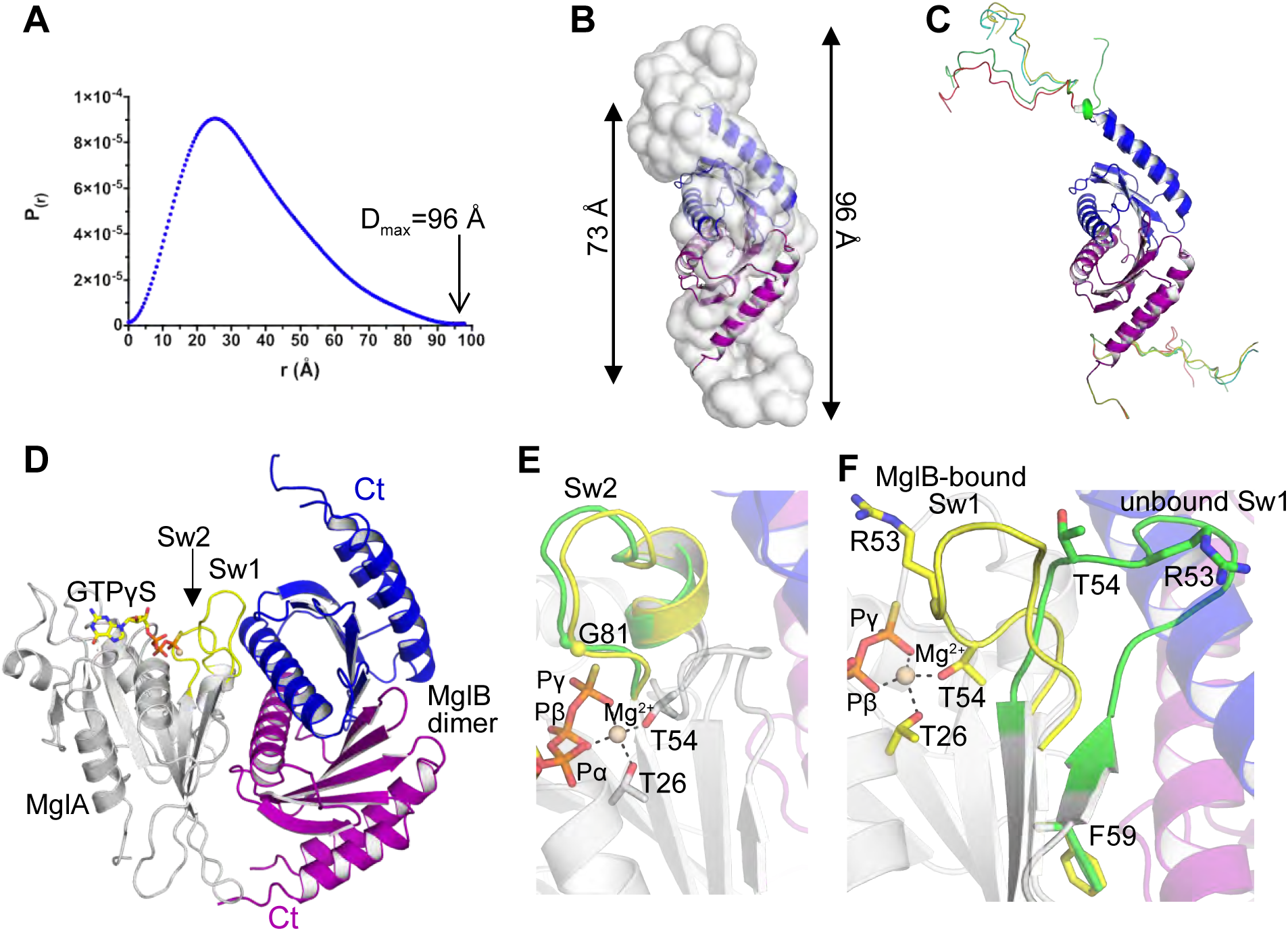
Crystallographic and solution structural analysis of MglB. **A-** SEC-SAXS distance distribution function P(r) of unbound MglB in solution. The D_max_ value is indicated. Complementary information on SAXS data collection and analysis is given in **Figures S1C-E**. **B-** Comparison of the crystallographic MglB dimer (in blue and purple) with the SAXS envelope calculated with GASBOR. The C-terminii are not visible in the crystal structure, but appear as extra volumes in the SAXS envelope (arrows). The Dmax values calculated from the crystal structure and the SAXS data are indicated. **C-** A composite model of the MglB dimer, in which the C-terminii is represented by ensemble structures calculated with MultiFoXS. **D-** Crystallographic structure of the MglA-GTPγS/MglB heterotrimer. MglB interacts with the switch 1 and with non-switch regions of MglA, forming an interface that is asymmetrical on the MglA side and symmetrical on the MglB dimer side. The position of the C-terminii of MglB are not visible in the crystal and are represented by dotted lines. **E-** Close-up view of the overlay of the switch 2 in unbound (green) and MglB-bound (yellow) MglA-GTPγS, showing that the conformation of the switch 2 is not remodelled by MglB. **F-** Close-up view of the overlay of the switch 1 in unbound (green) and MglB-bound (yellow) MglA-GTPγS, showing that MglB induces a lasso movement of the arginine finger loop that resolves steric conflicts and brings Arg53 and Thr54 into the nucleotide-binding site.

Next, we determined the crystal structure of the *M. xanthus* MglA-GTPγS/MglB complex. The complex contains a symmetrical MglB dimer and one MglA-GTPγS molecule (**Figure 2D**), as previously observed for *T. thermophilus* MglA-MglB complexes (Miertzschke et al., 2011). As in unbound MglB, the C-terminii of MglB are disordered in the MglA-MglB complex. Both MglB monomers interact through helix 2 with the switch 1 and switch 2 of MglA, which display active conformations that resemble those seen in unbound MglA-GTPγS **(Figures 2E and 2F)**, with additional contacts established outside the switch regions by the second monomer. The MglB dimer does not insert any element in the nucleotide-binding site and does not interact with GTPγS directly. Instead, the first monomer, which would clash with the arginine finger loop in unbound MglA-GTPγS resolves the conflict by displacing the arginine finger loop by a large lasso movement (11 Å at Arg53) (**Figure 2F**). This rearrangement positions the arginine finger such as to reach into the active site through side chain rotation, and completes the canonical coordination of Mg^2+^ by the conserved switch 1 threonine **(Figure 2F).** Thus, MglB recognizes the active conformations of the switch regions and contributes to catalysis by reorganizing the arginine finger loop through steric conflict. The organization of the MglA-MglB complex is surprisingly similar to that of the complex of the Rab GTPase Ypt1 with the TRAPP complex, a yeast RabGEF whose central subunits have roadblock folds (**Figure S1G**) (Cai et al., 2008). Therefore, we tested whether MglB might also act as a GEF for MglA, by measuring nucleotide exchange using fluorescence kinetics. As shown in **Figure S1H**, while MglA undergoes measurable spontaneous GDP/GTP exchange in the presence of Mg^2+^, this was not increased by MglB, indicating that MglB does not function as a GEF under these conditions.

### A positive feedback loop converts MglA-GTP from an MglB-resistant form to an MglB-sensitive form

The identification of a mixed MglA state, which has structural features to bind GTP but does not have the active switch 1 conformation seen in the MglA/MglB complex, raises the question of whether it plays a role in MglB-stimulated GTP hydrolysis. To get insight into this issue, we characterized the kinetics of MglB-stimulated GTP hydrolysis stimulated by fluorescence, using an engineered bacterial phosphate-binding protein (Brune et al., 1998, Mishra et al., 2013). Potent GAP activity was measured with this assay, which depended on the presence of MglB and GTP **(Figure S2A).** GTP hydrolysis kinetics measured over a range of MglB concentrations yielded a *k*_cat_/*K*_M_ of 2.1×10^3^ M^−1^ s^−1^ **(Figures 3A and S2B)**. While carrying out these experiments, we made the intriguing observation that MglA-GTP samples became gradually less sensitive to MglB over time when incubated at 25 °C and were resistant to GTP hydrolysis after about one hour (**Figure 3B**). In contrast, MglB remained equally active over time, indicating that the effect is entirely comprised in MglA. This effect was not due to unfolding, as checked by circular dichroism (**Figure S2C**). Analysis of the nucleotide content in MglB-insensitive samples showed that the major nucleotide bound to MglA was GTP, indicating that MglA-GTP did not convert spontaneously to MglA-GDP-Pi **(Figures S2D and S2E)**.

**Figure 3.**
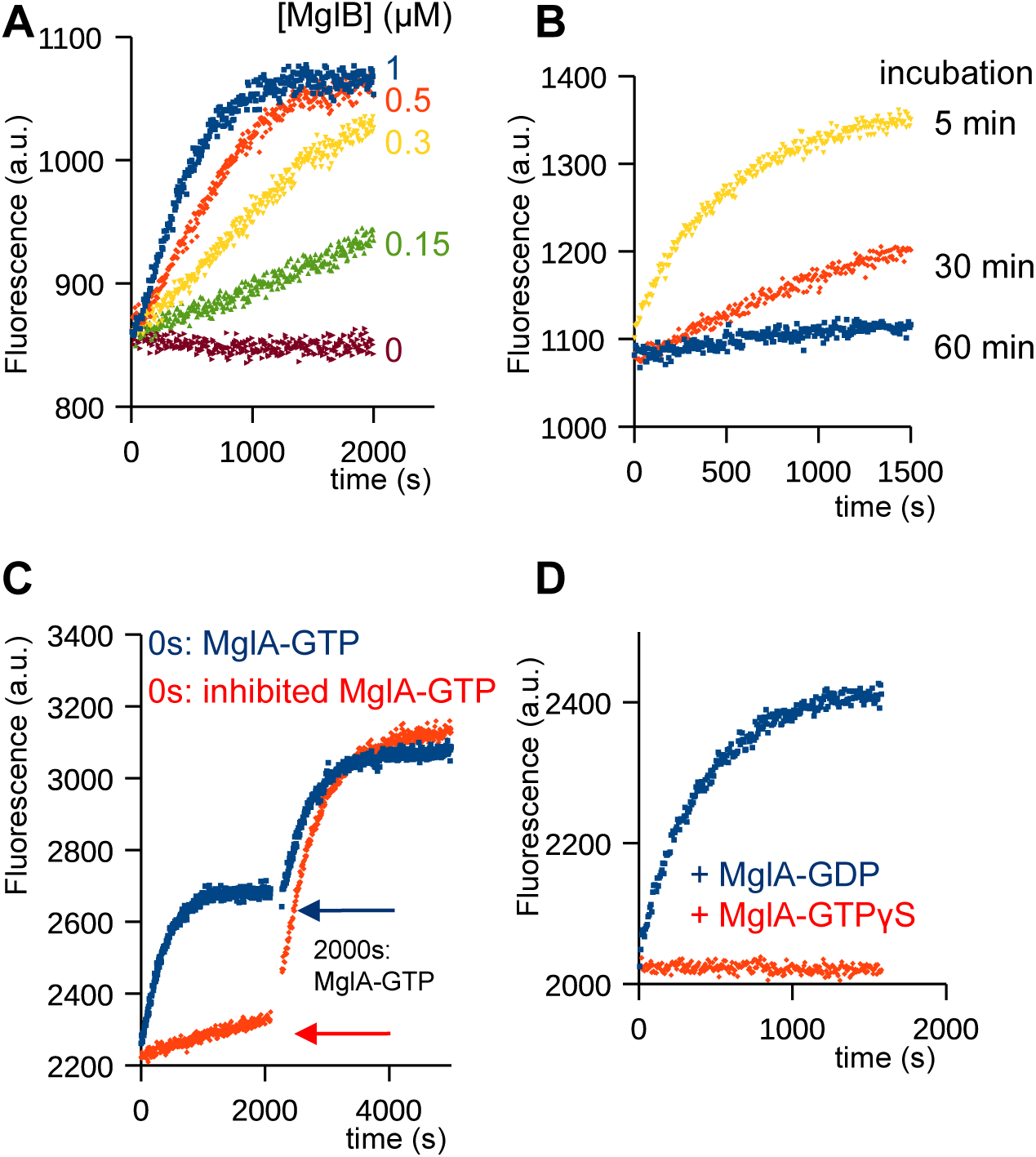
Inactivation of MglA by MglB is regulated by a positive feedback loop. **A-** Kinetics of GTP hydrolysis stimulated by MglB. Kinetics were monitored by fluorescence using a reagentless phosphate-binding assay at a fixed MglA concentration (1 µM) and increasing concentrations of MglB as indicated. **B-** Time-dependent desensitization of MglA to MglB-stimulated GTP hydrolysis. MglA-GTP was incubated at 25 °C for increasing periods of time as indicated, before GTP hydrolysis was initiated by addition of MglB. **C-** Desensitization of MglA from MglB is reversed by addition of MglB-sensitive MglA-GTP. MglB was added to MglB-insensitive (in orange) or MglB-sensitive (in blue) MglA-GTP and GTP hydrolysis was monitored for the duration needed for the MglB-sensitive sample to reach the plateau. Then, the same amount of MglB-sensitive MglA-GTP was added to both samples (arrow). Note that both kinetics curves reach the same plateau, indicating that all MglA-GTP has been converted to MglA-GDP in each experiments. **D-** MglA-GDP, but not MglA-GTPγS, resensitize MglA-GTP to MglB. MglA-GTP was incubated at 25 °C for 30 minutes, then MglB and 1µM of either MglA-GDP or MglA-GTPγS was added.

Next, we investigated whether resistance to MglB could be reverted. Remarkably, MglB-resistant MglA-GTP could be fully re-sensitized to MglB by addition of MglB-sensitive MglA-GTP (**Figure 3C**). No re-sensitization was observed by addition of GTP alone or of MglA-GTPγS, which cannot be converted to MglA-GDP, while addition of MglA-GDP fully re-sensitized MglA-GTP **(Figure 3D)**. These observations indicate that the re-sensitizing species is MglA-GDP, which in the experiment shown in **Figure 3C** is produced by MglB. We conclude from these experiments that MglA-GTP exists in two functional states: one which is sensitive to MglB and one which is insensitive to MglB and can be re-sensitized by MglA-GDP produced by MglB through a positive feedback loop.

### A three-state GTPase switch controls directed motility in *M. xanthus*

Our structural and biochemical analyses each identify an atypical MglA species, which has a mixed conformation that can accommodate GTP but is insensitive to MglB. To assess the existence and functional significance of this species *in vivo*, we designed a structure-based MglB mutant able to diffuse freely in the cytosol, such that it could reach all MglA-GTP in the cell. The MglB dimer features an extended, positively charged, convex tract located opposite to the MglA-binding site **(Figure 4A**) which is conserved across bacterial species **(Figure S1I)**, hence appears well-suited to support intermolecular interactions that determine MglB segregation at the lagging pole. We mutated K14, K120 and R115 in this tract into alanines (MglB^3M^) **(Figure 4A)**, resulting in the removal of 6 positively charged residues from the surface of the dimer. MglB^3M^ was fully efficient at inactivating MglA *in vitro*, indicating that the tract is not involved in GTP hydrolysis **(Figure S3A).** *In vivo*, MglB^3M^ was stably expressed (**Figure S3B**). MglB^3M^ fused to the Neon-Green protein displayed a conspicuous diffuse pattern that contrasted with the mostly unipolar distribution of MglB^WT^ **(Figure 4B)**, thus identifying the tract as a critical determinant of MglB segregation at the lagging pole. Remarkably, most cells expressing MglB^3M^ were non-motile, thus affecting cells more severely than an *mglB* deletion mutant **(Figures 4C and S3C)**, suggesting that MglB^3M^ blocks the function of MglA. To confirm this hypothesis, we took advantage of an activating mutation at Gln82 in the switch2 (Q82L) which renders MglA insensitive to inactivation by MglB (Zhang et al., 2010, Miertzschke et al., 2011). MglA^Q82L^ impairs disassembly of the motility complex leading to pendulum movements. The frequency these movements is increased by deletion of MglB, which is explained by the formation of unproductive MglA^Q82L^-MglB complexes that remove MglA from the motility complex and therefore block its assembly, an effect that cannot occur in the *ΔmglB* background (Treuner-Lange et al., 2015). Pendulum movements induced by MglA^Q82L^ were also blocked by MglB^3M^ (**Figure 4D**), confirming that MglB^3M^ acts through MglA *in vivo*.

**Figure 4.**
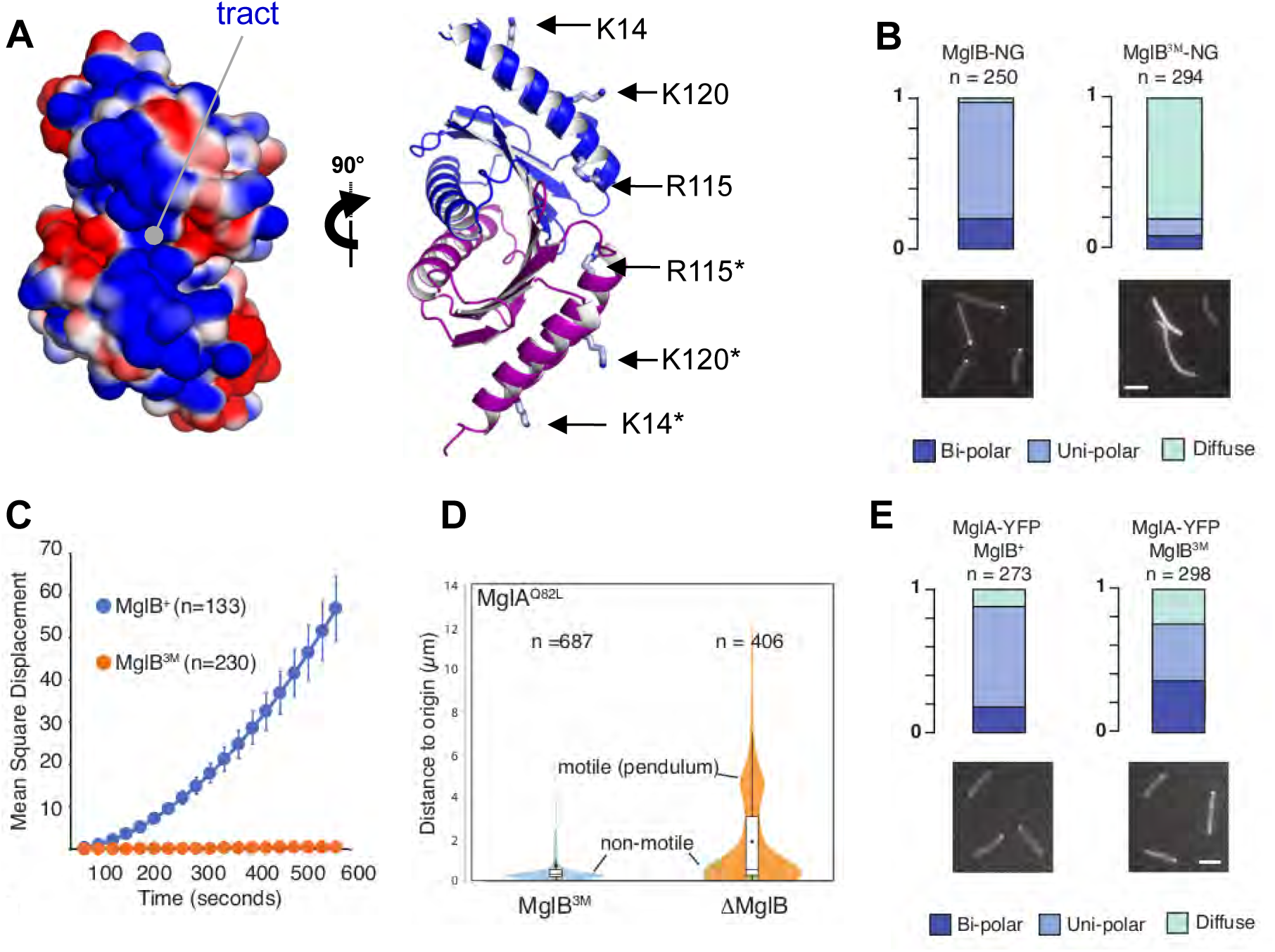
The MglB positively charged tract determines the polar localization of MglB. **A-** MglB features a positively charged tract opposite to the MglA-binding site. The left panel shows electrostatic potential calculated with APBS (Baker et al., 2001). The right panel shows the residues in this tract mutated in MglA^3M^. The two views are rotated by 90 degrees. **B-** MglB^3M^-NG has a diffuse localization. Localizations of MglB-NG and MglB^3M^-NG are shown. **C-** The expression of MglB^3M^ blocks motility. Motility is measured as the Mean Square Displacement (MSD) of single cells on agar. The average MSD of n trajectories and associated standard error of the mean are shown. **D-** MglB^3M^ blocks the action of a GTP-locked MglA mutant. The motility of MglA^Q82L^-expressing cells was measured in *mglB* deletion or in *mglB*^3M^-expressing strains. For each cell, motility is measured as the distance traveled from the origin for a time period of 45 min. Note that MglB^3M^-expressing cells are mostly non-motile while MglB^WT^-expressing cells show two distinct populations, one non-motile and one motile with reduced traveled distance to origin due to previously described pendulum movements. **E-** MglA-YFP retains polar localization in MglB^3M^-expressing cells. Localization of MglA-YFP in cells expressing MglB or MglB^3M^ is shown.

The above experiments show that MglB^3M^ has the required functional features to investigate whether all MglA-GTP, which has a mostly polar localization, can be converted by MglB into MglA-GDP, which is characterized by a cytosolic distribution (Leonardy et al., 2010, Miertzschke et al., 2011, Treuner-Lange et al., 2015). Remarkably, in MglB^3M^-expressing cells, MglA-YFP was still able to localize at the poles, indicating that a significant fraction of MglA-GTP is resistant to MglB-stimulated GTP hydrolysis even under conditions when the two proteins are no longer segregated from each other **(Figures 4E and S3D)**. Moreover, regulated polar switching of MglA was never observed (**Figure S3D**), thus explaining why cells fail to reverse and demonstrating that polar localization of MglB and both MglA-GTP species is required for the polarity switch.

Together, these observations provide several lines of evidence for the existence and functionality of the MglA-GTP* species *in vivo*: (i) Diffuse MglB^3M^ can reach the entire MglA-GTP pool, but an MglA-GTP subpopulation retains polar localization hence resists MglB-stimulated GTP hydrolysis; (ii) this polar MglA subpopulation cannot support motility, suggesting that it is structurally different from the MglA-GTP species that is able to engage productively with the motility machinery. These characteristics match those predicted for the mixed MglA-GTP* species observed in the crystal structure and for the MglB-resistant MglA-GTP* species identified in GAP kinetics, which altogether provides strong evidence that the MglA-GTP* species exists and serves a localization function *in vivo*.

## Discussion

In this study we investigated how the structural and biochemical features of the GDP/GTP switch of the small GTPase MglA contribute to molecular oscillations that control reversals in the gliding bacterium *M. xanthus*. We discovered that MglA has three functional states *in vitro* and *in vivo*, which were consistently observed 1) by X-ray crystallography, uncovering a mixed usage of active and inactive switch region conformations never seen in a small GTPase before, 2) by kinetics analysis, revealing that MglA-GTP converts from an MglB-sensitive to an MglB-resistant form (coined MglA-GTP*), in a manner that is reversed by a positive feedback loop operated by MglA-GDP, and 3) in live bacteria, using a diffusible MglB mutant, which reveals an MglB-resistant population (hence likely MglA-GTP*) that localizes to the cell poles. Our data suggests that the MglB-sensitive MglA-GTP species is represented by the MglA-GTPγS structure, in which both switch regions are in active conformations similar to those seen in the MglA/MglB complex (this study, (Miertzschke et al., 2011)). Accordingly, the MglA-GTP* species is best represented by the mixed MglA conformation, whose retracted switch 1 conformation cannot establish the same interactions with MglB as in the MglA/MglB complex, readily explaining its resistance to MglB-stimulated GTP hydrolysis. The mixed conformation does not feature Mg^2+^, a metal ion that has been shown to be crucial for GTP hydrolysis in other small GTPases (Zhang et al., 2000), possibly further impairing its ability to hydrolyze GTP. From a structural perspective, it is remarkable that the unique twisted retraction of the switch 1, previously thought to implement a classical inactive GDP-bound state, is in fact the signature of a 3-state GDP/GTP switch.

The existence of two distinct MglA-GTP species reveals that distinct functional MglA-GTP subpopulations operate in gliding bacteria: one which clusters stably at the leading edge to define polarity during movement, and one that associates with the motility complex and travels backwards to propel the bacterial. The transition between these subpopulations can potentially explain how MglA dissociates abruptly from the leading edge at the time of reversal (see below). Because of their structural differences (the conformation of the switch 1 region), MglA-GTP and MglA-GTP* have the potential to be discriminated by effectors and regulators, hence to define such subpopulations. Since MglA-GTP* is not sufficient for motility, it is predicted to bind to signaling components located at the leading pole. Likewise, MglA-GTP has active features that are predicted to allow it to recruit the motility complex, become transported to the lagging pole and be inactivated by MglB. Future studies are needed to determine whether MglA effectors and regulators bind in an exclusive manner to a specific MglA-GTP species and/or if they can bind either productively or unproductively depending on the MglA-GTP species. The regulation of this three-state GDP/GTP switch by a positive feedback loop operated by MglB and MglA-GDP potentially explains how MglA rapidly detaches from the leading pole to complete the reversal process (**Figure 5**). Our findings suggest that the MglA-GDP pool generated by inactivation of MglA-GTP during motility is the mobilization messenger that converts polar MglA-GTP* into diffusible MglA-GTP, allowing it to reach the lagging pole to be inactivated by MglB during reversals. This positive feedback loop would explain the fast non-linear dynamics of MglA re-localization to the lagging cell pole when reversals are provoked (Guzzo et al., 2018). Further work is needed to prove that the feedback loop operates *in vivo*, but the observation that an MglA-YFP population remains immobilized at the cell pole when MglB^3M^ is expressed strongly suggests that the MglA-GDP/MglA-GTP*/MglA-GTP cycle is required for functional MglA oscillations. Interestingly, the mixed MglA structure and MglA-GDP interact in the crystal through their retracted switch 1 (**Figure S2F**), suggesting that MglA-GDP form heterodimers with MglA-GTP* to facilitate their conversion into MglA-GTP.

**Figure 5:**
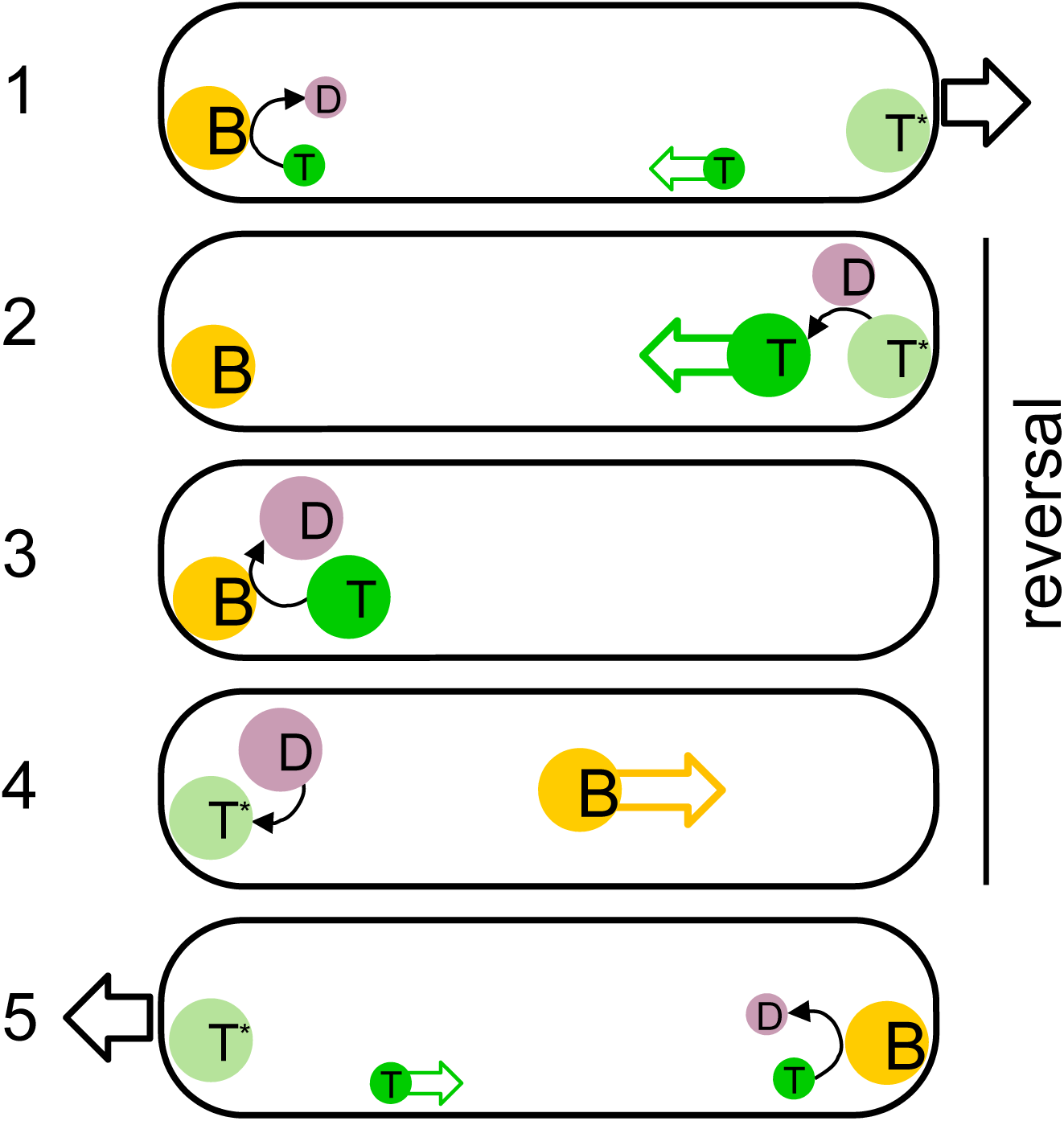
Model of the three-state MglA switch. 1-The MglA-GTP cluster located at the front pole of the gliding bacteria is composed of a major species stably associated to the pole which is resistant to MglB (MglA-GTP*, T* in kaki) and a minor diffusible species (MglA-GTP, T, in green) that can reach the lagging pole to be inactivated into MglA-GDP (D in mauve) by MglB (B, in orange). 2-During a reversal event, MglA-GDP is released in the cytosol to reach the leading pole where it converts all polar MglA-GTP* into diffusible MglA-GTP, which detaches rapidly from the pole. 3-MglA-GTP is rapidly inactivated into MglA-GDP by MglB at the lagging pole. 4-MglA-GDP is reloaded with GTP by an unknown GEF activity to regenerate MglB-resistant MglA-GTP*, and MglB relocates to the opposite pole. 5-A new leading pole is established, resuming movement in the opposite direction. The direction of movement at steps 1 and 5 is shown by a black arrow, the reversal event spanning steps 2-3-4 is indicated by a vertical line.

Finally, our study provides insight into the determinants of MglB clustering at the lagging pole. First, it identifies a conserved, positively charged surface as the major binding site for polar segregation of MglB. This region is located opposite to the MglA-binding site, suggesting that interactions that determine MglB localization are not mutually exclusive to its inactivation of MglA. Given the marked electrostatic character of the identified epitope, we asked whether MglB might interact directly with the membrane using liposomes enriched in cardiolipin, a negatively charged lipid enriched at bacterial poles (reviewed in (Romantsov et al., 2009)), using co-sedimentation. However, MglB^WT^ bound only weakly to liposomes, and binding was not measurably affected in MglB^3M^ (**Figure S3E**). We cannot exclude that the convex shape of the surface would be selective for membranes with negative curvature, which cannot be recapitulated by liposome experiments. Alternatively or in coordination, MglB localization to the cell pole may require protein partners, which remain to be identified. Second, we identified the C-terminii of the MglB dimer as intrinsically disordered regions, which considerably enlarge the volume occupied by MglB. Intrinsically disordered regions are increasingly recognized as providing polymer-like behaviors that support various functional features, such as recognition of ligands through multiple low affinity interactions (Milles et al., 2015), determination of optimal protein densities at membrane surfaces (Schmid et al., 2016) or sensing of membrane curvature (Zeno et al., 2018). The disordered C-terminii may thus endow MglB with such polymer-like features, and modulate thereby its segregation at the lagging pole. The detailed contributions of the positively charged surface and the intrinsically disordered C-terminii of MglB will have to be investigated.

The full understanding of the three-state GTPase switch will still require that important remaining issues are elucidated. One of them is the mechanism that controls the mobilization of the MglA-GDP pool at the onset of a reversal. In one mechanism, MglA-GDP could be generated rapidly by MglB following FrzX signaling. Alternatively, it may be actively retained at the lagging pole and protected from reloading GTP, until a signal releases it into the cytosol. Another important question will be to elucidate how MglA-GDP generated during the reversal is rapidly reloaded with GTP to complete the cycle. It is tempting to speculate that a GEF activity is upregulated during reversals, such as to convert MglA-GDP into MglA-GTP* to reconstitute the polarity axis. This atypical GEF activity may protect MglA from futile inactivation by MglB and contribute to MglB relocalization. In conclusion, we propose that the existence of three distinct MglA states is a key ingredient to enable MglB, RomR and MglA to communicate at a distance by means of a diffusible messenger, thereby implementing the spatial component of the relaxation oscillator switch. Given that small GTPases are involved in polarity control, it is likely that such regulation also operates in a number of other biological systems.

## Methods

### Protein cloning, expression and purification

All proteins were cloned in a pET28 plasmid. The MglA construct contains a non-cleavable 6xHis-tag at the C-terminus. Two MglB constructs were used. One construct carries a 34-residue tag at the N-terminus including a 6xHis tag and a thrombin cleavage site and was used for the crystal structure of unbound MglB (MglB-1). A second MglB construct carries a 6xHis tag followed by a TEV protease cleavage site at the N-terminus was used for all other structural and biochemical experiments (MglB-2). The MglB^K14A/R115A/K120A^ triple mutant (MglB^3M^) was obtained by inserting each mutation using a specific oligonucleotide starting from the MglB-2 plasmid.

All proteins were expressed in *E. coli* BL21(DE3) strain grown in LB medium. Overexpression was induced by 1 mM IPTG overnight at 20 °C. Bacteria were pelleted by centrifugation, resuspended in lysis buffer (Tris 20 mM pH 7.5, 300 mM NaCl, 20 mM imidazole, 5 mM DTT) complemented with benzonase and an anti-protease inhibitor cocktail and disrupted in a pressure cell homogeniser at 20000 psi and centrifuged. Cleared lysates were loaded onto a nickel affinity column (HisTrap FF, GE Healthcare) mounted on an Akta chromatography system. Proteins were eluted by a gradient of imidazole (20 to 500 mM). Purification of MglA was completed by gel filtration chromatography on a Superdex75 10/300 column (GE Healthcare) equilibrated with a buffer containing Tris 20 mM pH 8, 50 mM NaCl and 10 mM MgCl_2_. Purification of MglB-1 was completed by gel filtration chromatography on a Superdex75 16/600 column equilibrated with a buffer containing 50 mM Tris pH 8, 100 mM NaCl, 1 mM MgCl_2_. The 6xHis tag of MglB-2 was cleaved by incubation with the TEV protease overnight at 4°C (1:20 w/w ratio). The cleaved tag was removed by a second Ni^2+^ affinity step, then purification was completed by gel filtration chromatography on a Superdex75 10/300 column equilibrated with a buffer containing 20 mM Tris pH 8, 50 mM NaCl and 10 mM MgCl_2_. MglB^3M^ was purified as MglB-2. All proteins were pure over 90 % as judged by SDS-PAGE, concentrated to 2 mg/mL and stored at −20 °C.

Homogeneous loading of MglA with GDP, GTP, mant-GTP or GTPγS was achieved by incubation with a 20x molar excess of either nucleotide for 30 minutes at room temperature in a buffer containing 10 mM MgCl_2_, then excess nucleotide were removed by gel filtration using a Superdex75 column. We note that MglA-GDP and MglA-GTP elute at different volumes, which improves their chromatographic separation.

### Crystallization and X-ray diffraction data collection and processing

#### MglA-GDP

Initial crystallization conditions were identified with a sparse matrix screen (MPD Suite, Qiagen) by vapor diffusion in sitting drops on a Cartesian crystallization robot. After optimization, diffracting crystals were obtained in 0.2 M sodium tartrate and 40 % MPD. Crystals were cryo-protected with 10% glycerol and flash frozen before data collection at beamline PROXIMA 1 (SOLEIL synchrotron, Gif-sur-Yvette, France). Diffraction data were processed and scaled using the XDS package (Kabsch, 2010).

#### MglA-GTPγS

MglA-GTPγS was supplemented with a 5x molar excess of GTPγS before crystallization. Initial conditions were obtained with the PEGs II Suite screen (Qiagen) by vapor diffusion in sitting drops at 18°C using a Mosquito robot (TTP Labtech). Diffracting crystals were obtained in 0.2 M ammonium sulfate, 0.1 M sodium acetate pH 4.6, 12% (w/w) PEG 4000 in batch wells containing 1 μL of the sample with 1 μL of crystallization solution covered by paratone and kept at 4 °C. Crystals appeared after 5 to 6 days. Crystals were cryoprotected by transfer to the reservoir solution supplemented with 35% sucrose and flash frozen in liquid nitrogen. X-ray diffraction data were collected at beam line PROXIMA 2 A (SOLEIL synchrotron) and processed and scaled with the aP_scale module in autoPROC (Global Phasing Limited) (Vonrhein et al., 2011), using STARANISO to apply anisotropic correction to the data (Tickle et al., 2016).

#### MglA mixed form

A unique crystal was obtained with MglA-GDP after a few months using a sparse matrix screen (Classics suite, Qiagen) in 0.1 M HEPES pH 7.5 and 70% MPD, by hanging drops dispensed with a Cartesian crystallization robot. The crystal was cryo-protected with 10% glycerol and flash frozen before data collection at beamline PROXIMA 1 (SOLEIL synchrotron). Diffraction data were processed using XDS (Kabsch, 2010) and scaled with Aimless (Evans and Murshudov, 2013).

#### Unbound MglB

Initial crystallization conditions were identified using sparse matrix screens with a Cartesian robot and optimized to produce diffracting crystals. Two crystal forms were obtained, one in 20% isopropanol and 10% PEG 4000 which was used to derive a low resolution envelope (form 1), and one in 0.1 M MES pH 6, 0.15 M ammonium sulfate and 15 % (w/w) PEG 4000 which was used for structure determination (form 2). Crystals were cryo-protected with 10 % glycerol and flash frozen before data collection at beamline PROXIMA 1 at SOLEIL synchrotron. Diffraction data were processed and scaled using the XDS package (Kabsch, 2010).

#### MglA-GTPγS/MglB complex

The complex was prepared by incubating equimolar amounts of MglA and MglB with a 20x molar excess of GTPγS for 30 minutes at room temperature followed by purification by size-exclusion chromatography using a Superdex 75 column and concentration at 20 mg/mL with a 5x molar excess of GTPγS. Initial crystallization conditions were obtained with the Protein Complex Suite (Qiagen) screened by sitting drop-vapor diffusion at 18 °C using a Mosquito robot and were optimized in hanging drops by mixing 1 μL of the sample with 1 μL of the precipitant. Diffracting crystals grew in one week in 0.2 M HEPES pH 7 and 16% PEG 4000. Crystals were cryoprotected by transfer to a solution containing 0.2 M HEPES pH 7 and 35 % PEG 4000 and flash frozen in liquid nitrogen. X-ray diffraction data was collected at beam line ID30-B at the European Synchrotron Radiation Facility (Grenoble, France). Diffraction data were processed and scaled with autoPROC (Global Phasing Limited) (Vonrhein et al., 2011), using STARANISO to apply anisotropic correction to the data (Tickle et al., 2016).

### Structure determination and refinement

Structures of MglA-GDP, MglA-GTPγS and the MglA mixed form were solved by molecular replacement using Phaser (McCoy et al., 2007) from the PHENIX suite (Adams et al., 2010). The structure of *T. thermophilus* MglA-GDP (PDB 3T1O, (Miertzschke et al., 2011)) was used as a search model to solve the MglA-GDP structure. The refined structure of *M. xanthus* MglA-GDP was then used as a search model for subsequent molecular replacements. The structure of the MglA-GTPγS-MglB complex was solved by molecular replacement using the refined structures of MglB and MglA-GTPγS as search models.

No molecular replacement solution could be obtained for unbound MglB form 1 or form 2 crystals using *T. thermophilus* MglB as a search model. Alternatively, we used both crystals forms for molecular replacement with AMoRe (Navaza, 1994) followed by density modification (Cowtan, 2010, Kleywegt and Jones, 1994). First, molecular replacement was carried out with monoclinic (P2_1_) data (form 1, 15-4 Å) by using a monomeric multiple model (nine superimposed structures of *T thermophilus* MglB monomers from PDB entries 1J3W, 3T1R, 3T1S with pruned side-chains) and by applying rotational non-crystallographic symmetry (NCS) constraints based on self-rotation function peaks. The four monomeric models placed in the P2_1_ asymmetric unit appeared in an arrangement similar to the tetrameric structure of *T. thermophilus* MglB, though with imperfect 222 symmetry. Successive density modification including 4-fold NCS averaging and solvent flattening and phase extension to 3.3 Å provided us with electron density maps of mediocre quality, perhaps due to incompleteness of data though this was not further investigated, but sufficient to distinguish and carve out the average density of a monomeric unit. This density was then used as a molecular replacement probe to phase the P2_1_2_1_2_1_ orthorhombic data (form 2 crystal). Twenty copies of the monomer, corresponding to five tetramers, were placed in the P2_1_2_1_2_1_ asymmetric unit using data in the 15-4 Å range and by applying NCS constraints based on self-rotation function peaks. Successive density modifications by 20-fold NCS averaging and solvent flattening and phase extension to 2.4 Å yielded good quality and interpretable density maps for all chains.

Refinements were performed by multiple rounds of refinements with PHENIX (Afonine et al., 2012) and manual fitting in Coot (Emsley et al., 2010). Crystallographic statistics are given in **Table S1**. Structural data have been deposited with the Protein Data Bank with accession codes given in **Table 1**.

### Nucleotide exchange and GAP kinetics assays

Nucleotide exchange kinetics were measured by monitoring the decrease in fluorescence following mant-GDP dissociation and replacement by GTP added in 5-fold molar excess (λ_Ex_ = 360 nm, λ_Em_ = 440 nm).

GAP kinetics were measured by fluorescence, using an engineered bacterial phosphate-binding protein (PBP) (kind gift of Martin Webb, UK) to detect inorganic phosphate produced by GTP hydrolysis (λ_Ex_ = 430 nm, λ_Em_ = 465 nm). After expression and purification, PBP was labeled with N-[2-(1-maleimidyl)ethyl]-7-(diethylamino)coumarin-3-carboxamide (MDCC) as described (Brune et al., 1998). GAP experiments were performed at 25 °C using 5 μm (**Figures 3A and 3B**) or 10 µM (**Figures 3C and 3D**) of the PBP sensor, 1 μm of MglA and a range of MglB concentrations as indicated. Fluorescence measurements were done in a FlexStation Multi-Mode Microplate reader. *k_obs_* values were determined from a mono-exponential fit. k_cat_/K_m_ were determined by linear regression from *k_obs_* values measured over a range of GAP concentrations following the Michaelis–Menten model. All experiments were done in triplicate.

## SEC-MALS

For size exclusion chromatography coupled to multiangle light scattering (SEC-MALS) analyses, MglB was prepared at 3 mg/mL in a buffer containing 20 mM Tris pH 7.5, 50 mM NaCl, 10 mM MgCl_2_. The same buffer was used as the mobile phase for SEC using a Superdex 75 10/300 GL column on a Shimadzu HPLC. Multi-angle light scattering was detected with a MiniDAWN TREOS light scattering module and a refractometer Optilab T-rEX (Wyatt Technology).

### Circular dichroism

Measurements were performed at 25 °C using a Jobin-Yvon Marker IV high sensitivity dichrograph. The MglA sample was dialyzed in 20 mM phosphate buffer at pH 8 and kept at 4 °C. Measurements were recorded at 0, 10 and 30 minutes after placing the sample at 25 °C. Far-UV spectra were collected from 190 to 260 nm with 1 nm steps in a 0.1 cm path-length quartz cell. Three scans were averaged and corrected by subtracting a buffer spectrum.

### Nucleotide content determination

MglA was prepared in a GTP state as described in the purification section. Samples incubated for one hour on ice or at at 25 °C were denatured by addition of methanol at −20 °C for two hours and then centrifuged to remove the precipitated protein. The nucleotides in the soluble fractions were analyzed by ion exchange in a MonoQ column recording absorbance at 254 nm. Pure GTP and GDP diluted in the same buffer were analyzed as a reference.

### SAXS Data Collection, Analysis and Structural Modeling

MglB SAXS data was collected using the inline HPLC-coupled SAXS instrument at SWING beamline (SOLEIL Synchrotron, France). 600 μg MglB in a 40 µL volume (15 mg/ml) was injected into a size exclusion chromatography column (SEC-3 300 Å, Agilent Technologies, Inc.) equilibrated with elution buffer (20 mM Tris pH 8.0, 150 mM NaCl and 1mM DTT), prior to SAXS data acquisition. Data reduction to absolute units, frame averaging, and subtraction were done using the FOXTROT program (synchrotron SOLEIL). Frames corresponding to the high-intensity fractions of the peak and having constant radius of gyration (*R*_g_) were averaged. All SAXS data analyses were performed with programs from the ATSAS package (Petoukhov et al., 2012). R_g_ was evaluated using the data within the range of Guinier approximation sR_g_<1.3 and by the Guinier Wizard and Distance Distribution Wizard. The maximum distance D_max_ was estimated with PRIMUS and refined by trial and error with GNOM. The distance distribution functions P_(r)_ were calculated with GNOM. The dimensionless Kratky plot was calculated by plotting (qR_g_)^2^I_(q)_/I_(0)_ against qR_g_. The molecular weight was estimated by PRIMUS Molecular Weight wizard. The fit between scattering experimental amplitudes and amplitudes calculated from the crystal structure of the MglB dimer was calculated with CRYSOL. 10 independent *ab initio* models were calculated with GASBOR, using data at q=0.25 and imposing P2 symmetry, which were compared with SUPCOMB and clustered with DAMCLUST. The consensus model was represented by the lowest Normalized Spatial Discrepancy (NSD), which was determined with DAMSEL. The flexible C-terminal fragments in each MglB monomer were modelled with MultiFoXS (Schneidman-Duhovny et al., 2016). The resulting models were clustered into 4 ensembles, yielding an excellent fit to the experimental SAXS data. The SAXS data have been deposited with the SAXSDB database under the accession code SASDET9. SAXS statistics are given in **Table S2**.

### Liposome co-sedimentation assays

All lipids are natural lipids from Avanti Polar Lipids. Liposomes were prepared with 76% phosphatidylethanolamine (PE), 4.9% phosphatidylglycerol (PG), 9.3% cardiolipin (CL), 6.5% phosphatidylserine (PS), and 3% lysophosphatidylcholine (LPC) in 50 mM Tris buffer pH 7.5 with 220 mM sucrose. After 5 cycles of freezing and thawing, liposomes were extruded through a 0.2 μm polycarbonate filter. Sucrose was removed by dilution in a buffer containing 50 mM Tris pH 7.5 and 120 mM NaCl, followed by centrifugation at 100,000 rpm and resuspension in the same buffer. Co-sedimentation assays were performed by incubating proteins (1 μM) with liposomes (1 mM) at room temperature for 10 minutes, followed by centrifugation at 100,000 rpm for 20 minutes. Controls were prepared without liposomes. Supernatants were recovered and pellets were re-suspended in the original buffer volume. All samples were analyzed by SDS-PAGE. All experiments were done in triplicate.

### Bacterial strains, plasmids, growth conditions, and genetic constructs

Strains, plasmids and primers used for this study are listed in **Tables S3, S4 and S5**. In general, *M*. *xanthus* strains were grown at 32°C in CYE rich media as previously described (Bustamante et al., 2004). Plasmids were introduced in *M*. *xanthus* by electroporation. Complementation, expression of the fusion and mutant proteins were obtained by ectopic integration of the genes of interest at the Mx8-phage attachment site (Zhang et al., 2010) under the control of their own promoter in appropriate deletion backgrounds. *E*. *coli* cells were grown under standard laboratory conditions in Luria-Bertani broth supplemented with antibiotics, if necessary.

### Expression of MglB and MglB^3M^

MglB and MglB^3M^ were expressed in the mglB deletion strain. The pSWU19 *MglB^3M^* for complementation of the *mglB* deletion was constructed by amplifying the *mglB* coding sequence. The mutated fragment was cloned into the pSWU19 vector by the one-step sequence- and ligation-independent cloning (SLIC) method as described (Jeong et al., 2012).

### Contruction of MglB- and MglB^3M^-neon green fusions

MglB-NG and derivatives were expressed by complementing the *mglB* deletion mutant (Zhang et al., 2010) by integration of pSWU19 *mglB-ng/mglB^3M^-ng* at the Mx8 Phage attachment site. For this, the MglB/MglB^3M^ (using pET28 *MglB^3M^* as a template, **Table S4**) and mNeon-green encoding sequences were amplified by PCR and mixed for cloning into the pSWU19 vector by the SLIC method.

### Motility assays on plate

Soft-agar motility was performed as previously described (Bustamante et al., 2004). In general, cells were grown up to an OD between 0.4 to 0.8 and concentrated at OD = 5 then spotted (10μL) on CYE 0.5% (soft). Colonies were photographed after 48 H.

### Fluorescence imaging and fluorescence intensity measurements

For phase-contrast and fluorescence microscopy, cells from exponentially growing cultures were concentrated to an OD=2 by centrifugation of 1 mL of culture and resuspended in the corresponding volume of TPM buffer (10 mM Tris-HCl, pH 7.6, 8 mM MgSO_4_, and 1 mM KH_2_PO_4_). Then, a drop of 2 µL was deposited on a coverslip and covered with a 1.5 % agar pad with TPM buffer. Microscopic analysis was performed using an automated and inverted epifluorescence microscope TE2000-E-PFS (Nikon, France) with a 100×/1.4 DLL objective and a CoolSNAP HQ2 camera (Photometrics). All fluorescence images were acquired with a minimal exposure time to minimize bleaching and phototoxicity effects.

### Cell tracking

Image analysis was performed with MicrobeJ using a FIJI-based tracking procedure developed for bacteria (Ducret et al., 2016). Cells were detected with MicrobeJ by thresholding the phase-contrast images after stabilization. Cells were tracked using MicrobeJ on a minimum of 40 frames by calculating all object distances between two consecutive frames and selecting the nearest objects. The computed trajectories were systematically verified manually and, when errors were encountered, the trajectories were removed. Analysis of the trajectories, distance to origin and MSD calculations were performed with MicrobeJ. For each strain, at least two biological replicates acquired independently were analyzed.

### Cluster counting

Image analysis was performed with FIJI with the cell counter plugin and under MicrobeJ with the Maxima detection system. For cluster detection, all fluorescence images were acquired with a 1 s exposure time for optimal signal-noise ratio.

### Western blots

Samples were grown at 32°C in CYE medium to an optical density (OD) at 600 nm (OD600) of 0.4 to 1 a volume of culture equivalent to 1mL was centrifuged for 5 min at 7000 rpm. The pellet corresponding to whole cells was resuspended in SDS-PAGE loading buffer containing β-mercaptoethanol to an 10 OD600 units and heated for 10 min at 99°C. Proteins samples equivalent 1 OD600 unit were separated by SDS-PAGE. Electrophoresis was performed at 180V for 50 min at room temperature using 10% SDS-polyacrylamide gel. For Western blotting, proteins were transferred from gels onto nitrocellulose membranes. The membranes were blocked during 1h at room temperature in Tris-buffered saline (pH 7.6), 5% milk, 0.2% Tween 20 (for MglA) or in Tris-buffered saline (pH 7.6), 2% milk, 0.2% Tween 20 (for MglB) and incubated with primary antibodies directed against MglA (dilution at **1:**5000) or MglB (dilution at **1:**2500) in blocking buffer overnight at 4°C. After two washings of 5 min with Tris-buffered saline (pH 7.6), 0.2% Tween 20 membrane were incubated with Goat Anti-Rabbit IgG (H + L)-HRP Conjugate (#1706515 biorad) in the respective blockings buffers. The peroxidase reaction was developed by chemiluminescence (SuperSignal™ West Pico Chemiluminescent Substrate #34080 Thermo Scientific™) scanned and analyzed with ImageQuant LAS 4000 and TL analysis software (GE Healthcare life sciences).

## Acknowledgements

This work was supported by grants from the ANR to J.C. and T.M. (ANR-15-CE13-0006) and to J.C. (ANR-14-CE09-0028). We are grateful to Armelle Vigouroux (LEBS, CNRS, Gif-sur-Yvette) for preliminary crystallization experiments. We thank the scientific staff at X-ray crystallography beam lines PROXIMA1 and PROXIMA 2 and SAXS beamline SWING (SOLEIL synchrotron, Gif-sur-Yvette, France) and X-ray crystallography ID-29 (ESRF synchrotron, Grenoble, France) for making the beamlines available to us and for expert advice.

---

**Table S1.** Crystallographic data and refinement statistics. Numbers in parentheses refer to the highest resolution shell.

| | MglA-GDP | MglA-GTPγS | MglA-GTPγS-MglB | MglA-GDP-Pi/MglA-GDP (mixed form) | MglB |
| --- | --- | --- | --- | --- | --- |
| **Data collection** |  |  |  |  |  |
| X-ray beamline source | PX-1 (SOLEIL) | PX-2a (SOLEIL) | ID-30b (ESRF) | PX-1 (SOLEIL) | PX-1 (SOLEIL) |
| Resolution range | 36.13-1.98 (2.05-1.98) | 47.91 - 1.28 (1.32- 1.28) | 44.22 - 2.8 (2.9 - 2.8) | 48.72 - 2.3 (2.38 - 2.3) | 45.41-2.39 (2.48-2.39) |
| Space group | P 2<sub>1</sub> 2<sub>1</sub> 2<sub>1</sub> | I 2 3 | P 6<sub>4</sub> | P 2<sub>1</sub> 2<sub>1</sub> 2 | P 2<sub>1</sub> 2<sub>1</sub> 2<sub>1</sub> |
| Unit cell<br>a, b, c (Å) | 54.02,<br>89.2,<br>118.55 | 117.35,<br>117.35,<br>117.35 | 135.10,<br>135.10,<br>60.38 | 88.6,<br>116.65,<br>49.11 | 119.78,<br>139.6,<br>179.42 |
| $\alpha, \beta, \gamma$ (°) | 90, 90, 90 | 90, 90, 90 | 90, 90, 120 | 90, 90, 90 | 90, 90, 90 |
| Multiplicity | 4.4 (4.1) | 6.8 (6.8) | 6.2 (5.9) | 5.3 (4.7) | 9.0 (8.9) |
| Completeness (%) | 98 (100) | 96.2 (63.2) | 69.5 (19.9) | 74.7 (20.5) | 73 (97) |
| I/$\sigma$(I) | 13.07 (0.60) | 14.01 (1.43) | 14.49 (1.90) | 8.29 (1.03) | 15.36 (1.10) |
| R-meas | 0.14 (1.28) | 0.07 (1.11) | 0.08 (1.09) | 0.18 (1.48) | 0.09 (2.04) |
| CC1/2 | 99.4 (74.4) | 99.9 (60.9) | 99.9 (65.8) | 99.3 (0.39) | 99.9 (49.4) |
| **Refinement** |  |  |  |  |  |
| Reflections used in refinement | 40036 (3634) | 67062 (4392) | 10923 (309) | 17471 (473) | 86633 (1124) |
| R-work / R-free | 21.4 / 23.8 | 16.0 / 17.73 | 20.53 / 25.25 | 20.88 / 26.79 | 26.36 / 28.61 |
| RMS deviations |  |  |  |  |  |
| Bonds (Å) | 0.014 | 0.006 | 0.002 | 0.002 | 0.012 |
| Angles (°) | 1.79 | 0.97 | 0.62 | 0.68 | 1.60 |
| Ramachandran favored (%) | 99 | 96.86 | 93.14 | 95.03 | 97 |
| Ramachandran outliers (%) | 0 | 0.00 | 1.37 | 0.79 | 0.57 |
| Average B-factor | 47.89 | 22.57 | 72.08 | 41.02 | 70.48 |
| solvent | 36.13 | 38.67 | 49.31 | 43.49 | 47.18 |

**Table S2.** SAXS data collection and analysis statistics.

|  |  |  |  |
| --- | --- | --- | --- |
| X-ray Beamline Source | SWING at SOLEIL Synchrotron, France |  |  |
| Wavelength | 1.03 Å |  |  |
| Detector | CCD (AVIEX170170) |  |  |
| Sample-detector distance | 1.8 m |  |  |
| Beam geometry | 0.4 x 0.1 mm |  |  |
| q range | 0.006-0.56 Å<sup>-1</sup> |  |  |
| Sample Injection | HPLC (Bio-SEC3 column) |  |  |
| Temperature | 15 °C |  |  |
| Exposure time | 2.0 s per frame |  |  |
| **Structural Parameters** |  |  |  |
| **From Guinier fit** |  | **From *P*<sub>(r)</sub>** |  |
| I<sub>(0)</sub> (cm<sup>-1</sup>) | 0.048±6e-05 | I<sub>(0)</sub> (cm<sup>-1</sup>) | 0.05 |
| R<sub>g</sub> (Å) | 27.61±0.33 | R<sub>g</sub> (Å) | 28.38 |
| qR<sub>g</sub> limits | 0.63-1.29 | D<sub>max</sub> (Å) | 96.0 |
|  |  | V<sub>P</sub> (nm<sup>3</sup>) | 56.4 |
| **Molecular mass determination** |  |  |  |
| MglB Monomer Theoretical Value | 17.3 kD |  |  |
| Qp (q<sub>max</sub>=0.25) | 40.71 kD |  |  |
| MoW (q<sub>max</sub>=0.45) | 36.51 kD |  |  |
| Vc (q<sub>max</sub>=0.30) | 37.80 kD |  |  |
| Bayesian inference | 37.70 kD |  |  |
| **Model Evaluation** |  |  |  |
| Crysol Fit to MglB Dimer* (Chi<sup>2</sup>) | 19.81 |  |  |
| MultiFoxS # (χ<sup>2</sup>) | 1.22 (e=4) |  |  |
| GASBOR (χ<sup>2</sup>) | 0.78 (NSD=1.04, n=5) |  |  |
| **SASDBD Accession Code (https://www.sasbdb.org)** |  | *SASDET9* |  |

Note: # Crystal structure of the MglB dimer with the two C-terminal peptides modeled as ensemble structures. NSD: Normalized Spatial Discrepancy

**Table S3.** Strains used in this study.

| Strain | Construction | Genotype | Source |
| --- | --- | --- | --- |
| TM155 | DZ2 ΔmglB | ΔmglB | (Zhang et al., 2010) |
| TM168 | TM155 mglB (pSWU19 mglB) | ΔmglB mglB | (Zhang et al., 2010) |
| TM1199 | TM155 mglB<sup>3M</sup> (pSWU19 mglB<sup>3M</sup>) | ΔmglB mglB<sup>3M</sup> | This work |
| TM1264 | TM155 mglB mglB-neon green (pSWU19 mglB-neon green) | ΔmglB mglB-neon green | This work |
| TM1265 | TM155 mglB mglB<sup>3M</sup>-neon green (pSWU19 mglB<sup>3M</sup>-neon green) | ΔmglB mglB<sup>3M</sup>-neon green | This work |
| TM1316 | ΔmglB mglA-YFP mglA (pSWU30 mglA) mglB (pSWU19 mglB) | ΔmglB mglA-yfp mglA mglB | This work |
| TM1319 | ΔmglB mglA-YFP mglA (pSWU30 mglA) mglB<sup>3M</sup> (pSWU19 mglB<sup>3M</sup>) | ΔmglB mglA-yfp mglA mglB<sup>3M</sup> | This work |

**Table S4.** Plasmids used in this study

| Plasmid | Relevant characteristics | Source or reference |
| --- | --- | --- |
| pSUW19 | KmR used to integrate genes ectopically at Mx8 att | (Leonardy et al., 2010) |
| pSUW19 MglB | pSWU19 with MglB under MglBpromoter | (Zhang et al., 2010) |
| pSUW19 mglB3M | pSWU19 with mglB<sup>3M</sup> under MglBpromoter | This work |
| pSUW30 mglB-neon green | pSWU30 with mglB-neon greenfusion under MglBpromoter | This work |
| pSUW19 mglB<sup>3M</sup>-neon green | pSWU19 with mglB<sup>3M</sup>-neon green under MglBpromoter | This work |

**Table S5.**
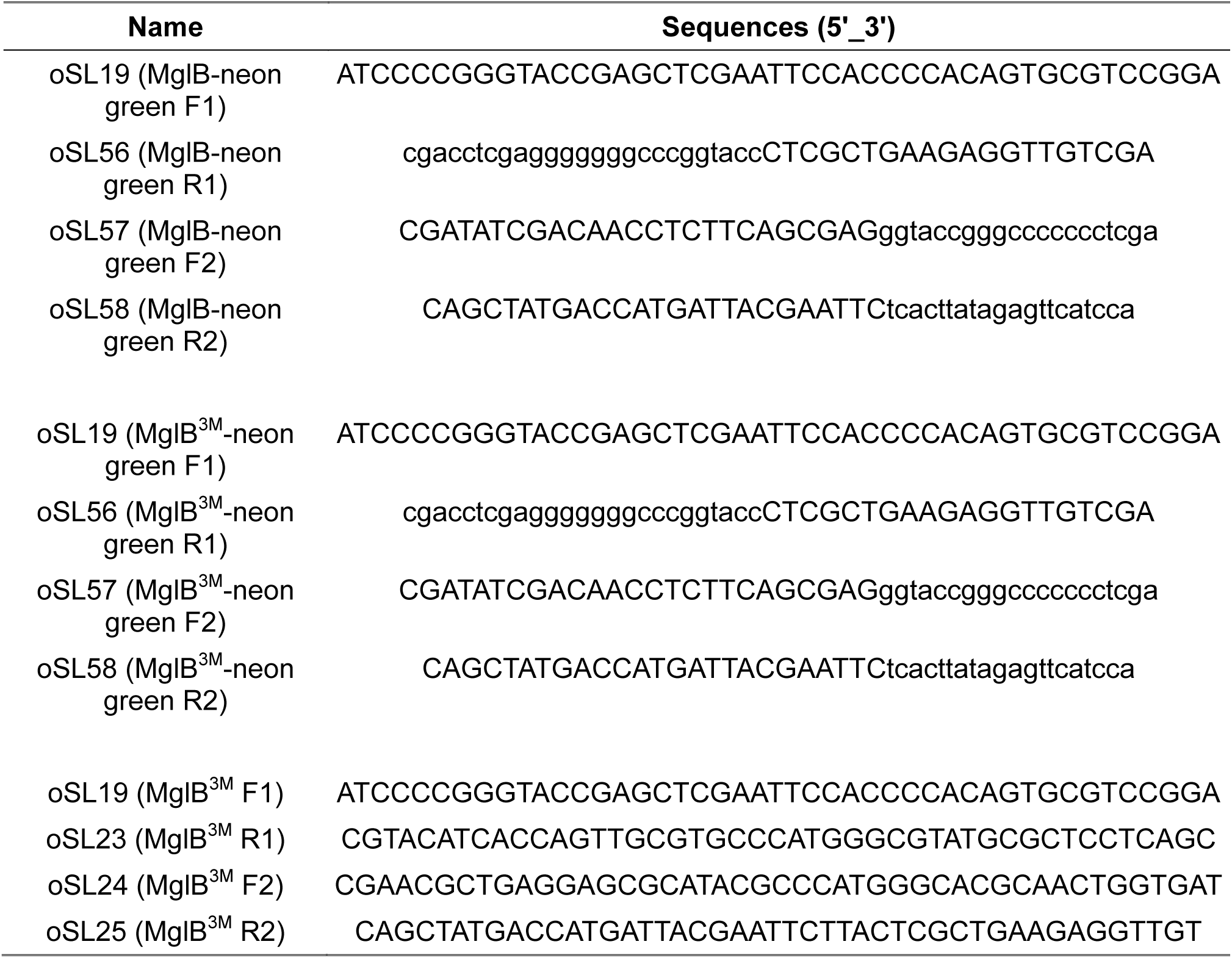
Primers used in this study.

**Figure S1.**
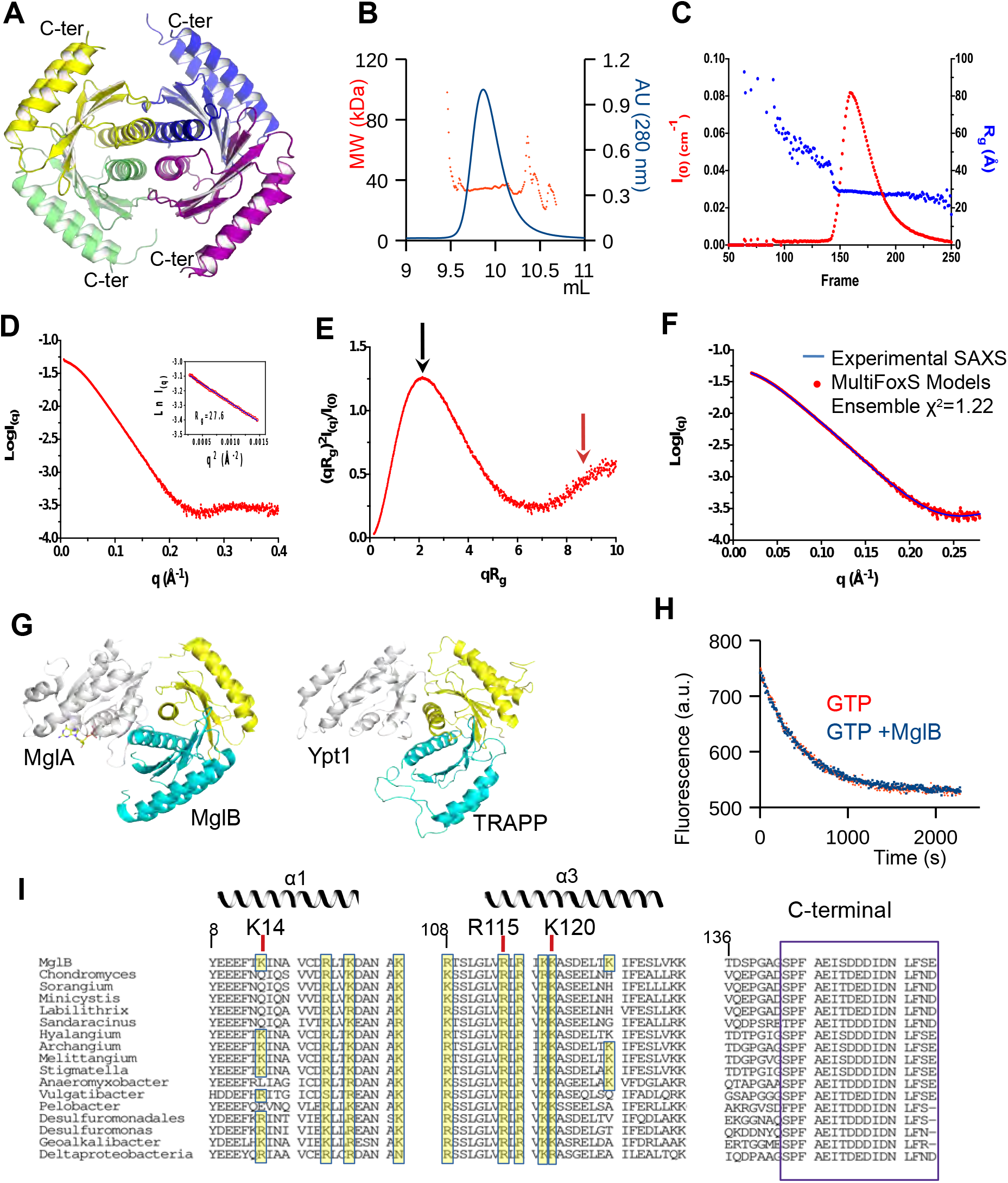
Crystal and solution structural analysis of MglB. Associated with Figure 2. **A-** Crystallographic structure of unbound MglB. The asymmetric unit contains 20 independent MglB molecules arranged as 5 tetramers as shown. The 26 residues in C-terminus are not visible in any of the monomers. The last visible residue is labelled C-ter. **B-** SEC-MALS analysis shows that MglB is an homogeneous dimer. **C-** Plot of I_(0)_ and R_g_ as a function of frame number for the SEC-SAXS analysis of MglB. Frames 168-187 were selected for data averaging. **D-** SEC-SAXS profile of MglB. The insert shows the Guinier plot (q_max_*Rg=1.19). The radius of gyration R_g_ is estimated from Guinier analysis. **E**-Dimensionless Kratky plot. The shifted peak value (black arrow) and the upwards shape at high qR_g_ (red arrow) are indicative of an elongated structure with flexible segments. **F-** The SAXS profile calculated from the MglB ensemble model with flexible C-terminii calculated with MultiFoXS (blue) has an excellent fit to the the experimental SAXS data (red). **G-** The MglA/MglB complex and the yeast YPT1-TRAPP complex ((Cai et al., 2008), PDB entry 3CUE) feature a common usage of the longin/roadblock dimer to bind their cognate small GTPase. **H-** MglA has measurable intrinsic nucleotide dissociation which is not further stimulated by MglB. Nucleotide exchange was monitored by following the replacement of mant-GDP by GTP by fluorescence (λ_Ex_ = 360 nm, λ_Em_ = 440 nm) in the presence (blue) or absence (orange) of MglB. MglA and MglB are at 1µM. Nucleotide dissociation is initiated by addition of 5 µM GTP. **I**-Sequence alignment of helices α2 and α3 and C-terminus of MglB from different bacteria species. Conserved positively charged residues in helices α2 and α3 are highlighted in yellow. Residues mutated in MglB^3M^ are labelled. Invariant or highly conserved residues in the flexible C-terminus are boxed.

**Figure S2.**
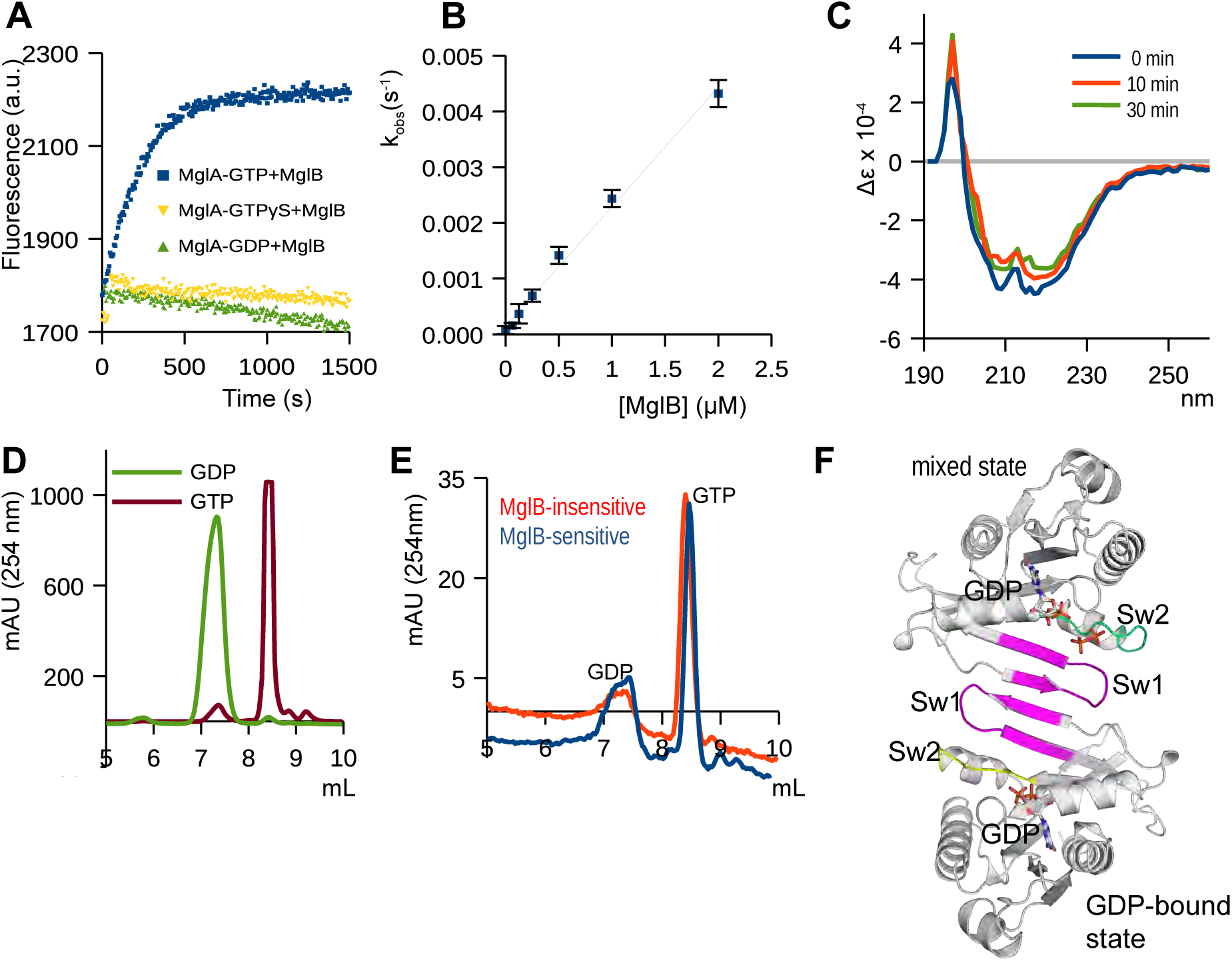
Characterization of the GAP activity of MglB. Associated with Figure 3. **A-** Specificity of MglB GAP kinetics measured by fluorescence with the reagentless phosphate-binding protein assay. Production of inorganic phosphate is observed with MglA-GTP (blue), but not with MglA loaded with GTPγS, a non-hydrolyzable GTP analog (red) or with MglA-GDP (green). **B-** Determination of k_cat_/K_m_ of GTP hydrolysis stimulated by MglB from a linear regression of *k_obs_* determined at different MglB concentrations. Individual *k_obs_* were determined by mono-exponential fitting. **C-** Circular dichroism spectra of MglA-GTP after incubation at 25 °C for 0 (blue), 10 (orange) and 30 (green) minutes. All spectra are representative of a folded protein. **D-** Elution volumes of GDP and GTP nucleotides monitored at 254 nm using MonoQ anion exchange chromatography, used as a reference for the experiments shown in Figure S2E. **E-** MglB-sensitive (incubated on ice, blue) and MglB-insensitive (incubated at 25 °C, red) MglA species contain GTP. MglA-GTP samples were denaturated by addition of methanol and centrifuged to remove precipitated protein before loading on the MonoQ column and analysis at 254 nm. **F-** MglA in the mixed state and MglA-GDP associate as an asymmetric dimer in the crystal through their switch 1 regions. The inactive switch 1 regions are shown in magenta, the active switch 2 in the mixed state is in green, the inactive switch 2 in MglA-GDP is in yellow.

**Figure S3.**
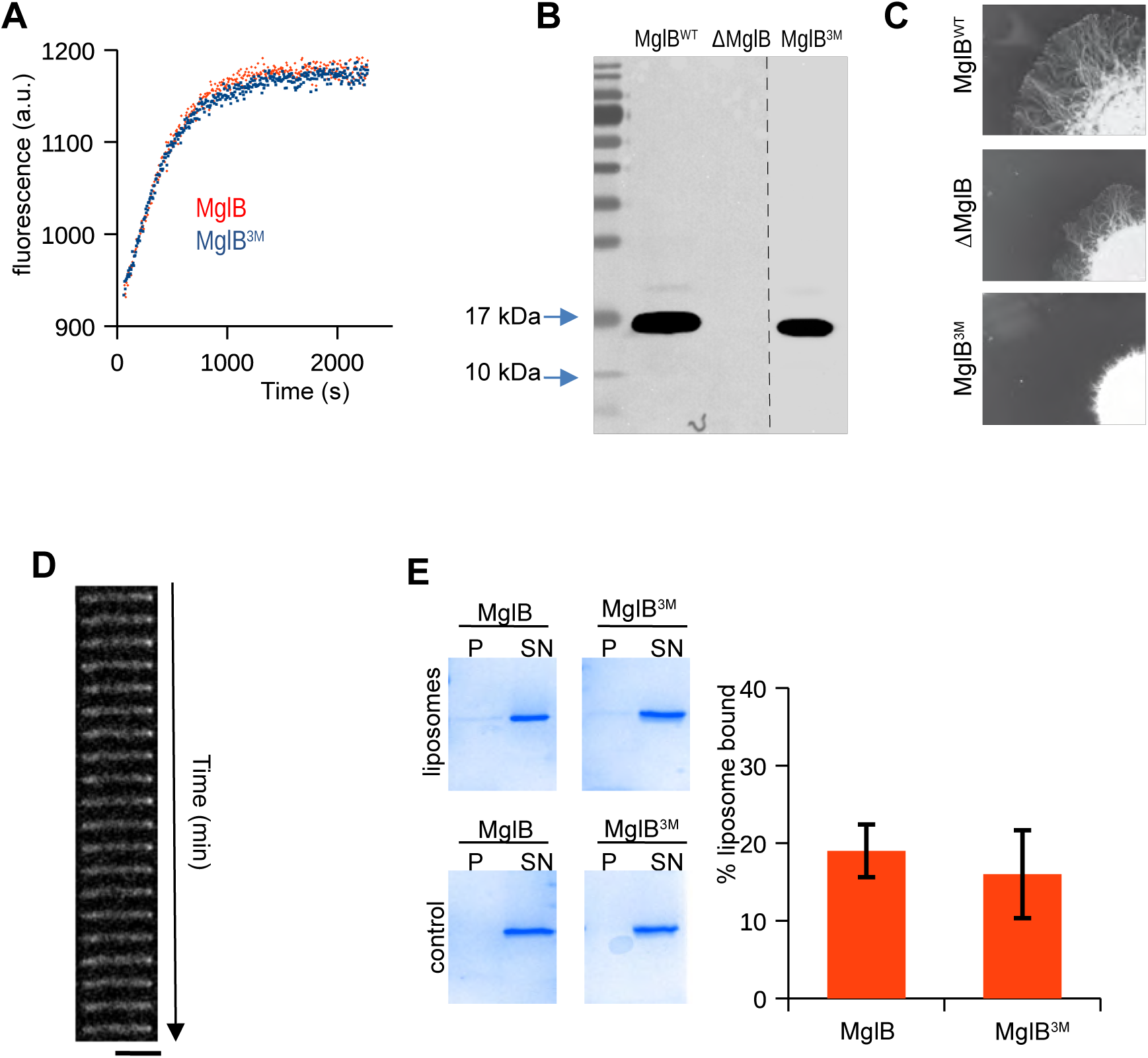
Analysis of the positively charged tract of MglB. Associated with Figure 4. **A-** The positively charged tract in MglB is not involved in the GAP activity. GAP kinetics were measured using the reagentless GAP assay. Wild-type MglB is in red, MglB^3M^ in blue. **B-** Stable expression of MglB^3M^. Shown is an anti-MglB Western blot. MglB and MglB^3M^ are stably expressed in an *mglB* deletion strain under the control of the *mglB* promoter inserted at the Mx8-phage attachment site. The dotted line shows that the MglB^3M^ blot was spliced closer to the MglB^WT^ and ΔMglB lanes for presentation purposes. **C-** Expression of MglB^3M^ leads to profound colony motility defects. Motility on 0.5% Agar is shown for each strain after 48H incubation at 32°C. Scale bar = 1 cm. **D-** MglA-YFP does not switch poles when MglB^3M^ is expressed. Shown is a time lapse example of MglA-YFP in an MglB^3M^-expressing cell. The fluorescence cluster is observed persistently at the same pole during the entire time lapse. 1 min time points are shown for a total duration of 20 min. scale bar = 2 µm. **E-** MglB binds weakly to cardiolipin-containing liposomes and this does not involve the positively charged tract. MglB-liposome interaction was analyzed by co-sedimentation. Pellets (P) containing liposome-bound proteins and supernatants (SN) were analyzed by SDS-PAGE. Controls experiments are carried out without liposomes. Right panel: quantification of the co-sedimention experiments.

## References

Adams, P. D., Afonine, P. V., BunkÓCZI, G., Chen, V. B., Davis, I. W., Echols, N., Headd, J. J., Hung, L.-W., Kapral, G. J., Grosse-Kunstleve, R. W., Mccoy, A. J., Moriarty, N. W., Oeffner, R., Read, R. J., Richardson, D. C., Richardson, J. S., Terwilliger, T. C. & Zwart, P. H. 2010. PHENIX: a comprehensive Python-based system for macromolecular structure solution. Acta crystallographica. Section D, Biological crystallography, 66, 213–221.

Afonine, P. V., Grosse-Kunstleve, R. W., Echols, N., Headd, J. J., Moriarty, N. W., Mustyakimov, M., Terwilliger, T. C., Urzhumtsev, A., Zwart, P. H. & Adams, P. D. 2012. Towards automated crystallographic structure refinement with phenix.refine. Acta crystallographica. Section D, Biological crystallography, 68, 352–67.

Baker, N. A., Sept, D., Joseph, S., Holst, M. J. & Mccammon, J. A. 2001. Electrostatics of nanosystems: application to microtubules and the ribosome. Proc Natl Acad Sci U S A, 98, 10037–41.

Brune, M., Hunter, J. L., Howell, S. A., Martin, S. R., Hazlett, T. L., Corrie, J. E. T. & Webb, M. R. 1998. Mechanism of inorganic phosphate interaction with phosphate binding protein from Escherichia coli. Biochemistry, 37, 10370–10380.

Bustamante, V. H., Martinez-FLORES, I., Vlamakis, H. C. & Zusman, D. R. 2004. Analysis of the Frz signal transduction system of Myxococcus xanthus shows the importance of the conserved C-terminal region of the cytoplasmic chemoreceptor FrzCD in sensing signals. Mol Microbiol, 53, 1501–13.

Cai, Y., Chin, H. F., Lazarova, D., Menon, S., Fu, C., Cai, H., Sclafani, A., Rodgers, D. W., De LA CRUZ, E. M., Ferro-NOVICK, S. & Reinisch, K. M. 2008. The structural basis for activation of the Rab Ypt1p by the TRAPP membrane-tethering complexes. Cell, 133, 1202–13.

Charest, P. G. & Firtel, R. A. 2007. Big roles for small GTPases in the control of directed cell movement. The Biochemical journal, 401, 377–390.

Cherfils, J. 2017. Encoding Allostery in mTOR Signaling: The Structure of the Rag GTPase/Ragulator Complex. Mol Cell, 68, 823–824.

Cherfils, J. & Zeghouf, M. 2011. Chronicles of the GTPase switch. Nat Chem Biol, 7, 493–5.

Cherfils, J. & Zeghouf, M. 2013. Regulation of small GTPases by GEFs, GAPs, and GDIs. Physiological reviews, 93, 269–309.

Cowtan, K. 2010. Recent developments in classical density modification. Acta Crystallographica Section D Biological Crystallography, 66, 470–478.

Ducret, A., Quardokus, E. M. & Brun, Y. V. 2016. MicrobeJ, a tool for high throughput bacterial cell detection and quantitative analysis. Nat Microbiol, 1, 16077.

Emsley, P., Lohkamp, B., Scott, W. G. & Cowtan, K. 2010. Features and development of Coot. Acta Crystallographica Section D: Biological Crystallography, 66, 486–501.

Evans, P. R. & Murshudov, G. N. 2013. How good are my data and what is the resolution? Acta Crystallographica Section D: Biological Crystallography, 69, 1204–1214.

Faure, L. M., Fiche, J. B., Espinosa, L., Ducret, A., Anantharaman, V., Luciano, J., Lhospice, S., Islam, S. T., Treguier, J., Sotes, M., Kuru, E., Van NIEUWENHZE, M. S., Brun, Y. V., Theodoly, O., Aravind, L., Nollmann, M. & Mignot, T. 2016. The mechanism of force transmission at bacterial focal adhesion complexes. Nature, 539, 530–535.

Guzzo, M., Murray, S. M., Martineau, E., Lhospice, S., Baronian, G., My, L., Zhang, Y., Espinosa, L., Vincentelli, R., Bratton, B. P., Shaevitz, J. W., Molle, V., Howard, M. & Mignot, T. 2018. A gated relaxation oscillator mediated by FrzX controls morphogenetic movements in Myxococcus xanthus. Nat Microbiol, 3, 948–959.

Islam, S. T. & Mignot, T. 2015. The mysterious nature of bacterial surface (gliding) motility: A focal adhesion-based mechanism in Myxococcus xanthus. Semin Cell Dev Biol, 46, 143–54.

Jeong, J. Y., Yim, H. S., Ryu, J. Y., Lee, H. S., Lee, J. H., Seen, D. S. & Kang, S. G. 2012. One-step sequence-and ligation-independent cloning as a rapid and versatile cloning method for functional genomics studies. Appl Environ Microbiol, 78, 5440–3.

Kabsch, W. 2010. Integration, scaling, space-group assignment and post-refinement. Acta Crystallographica Section D Biological Crystallography, 66, 133–144.

Keilberg, D., Wuichet, K., Drescher, F. & Sogaard-ANDERSEN, L. 2012. A response regulator interfaces between the Frz chemosensory system and the MglA/MglB GTPase/GAP module to regulate polarity in Myxococcus xanthus. PLoS Genet, 8, e1002951.

Kleywegt, G. J. & Jones, T. A. 1994. From First Map to Final Model, edited by S. Bailey, R. Hubbard & DA Waller, 59–66.

Leonardy, S., Miertzschke, M., Bulyha, I., Sperling, E., Wittinghofer, A. & SØGAARD-ANDERSEN, L. 2010. Regulation of dynamic polarity switching in bacteria by a Ras-like G-protein and its cognate GAP. The EMBO journal, 29, 2276–2289.

Mauriello, E. M. & Zusman, D. R. 2007. Polarity of motility systems in Myxococcus xanthus. Curr Opin Microbiol, 10, 624–9.

Mccoy, A. J., Grosse-KUNSTLEVE, R. W., Adams, P. D., Winn, M. D., Storoni, L. C. & Read, R. J. 2007. Phaser crystallographic software. Journal of Applied Crystallography, 40, 658–674.

Mercier, R. & Mignot, T. 2016. Regulations governing the multicellular lifestyle of Myxococcus xanthus. Curr Opin Microbiol, 34, 104–110.

Miertzschke, M., Koerner, C., Vetter, I. R., Keilberg, D., Hot, E., Leonardy, S., SØGAARD-ANDERSEN, L. & Wittinghofer, A. 2011. Structural analysis of the Ras-like G protein MglA and its cognate GAP MglB and implications for bacterial polarity. The EMBO Journal, 30, 4185–4197.

Milles, S., Mercadante, D., Aramburu, I. V., Jensen, M. R., Banterle, N., Koehler, C., Tyagi, S., Clarke, J., Shammas, S. L., Blackledge, M., Grater, F. & Lemke, E. A. 2015. Plasticity of an ultrafast interaction between nucleoporins and nuclear transport receptors. Cell, 163, 734–45.

Mishra, A. K., Del CAMPO, C. M., Collins, R. E., Roy, C. R. & Lambright, D. G. 2013. The Legionella pneumophila GTPase activating protein LepB accelerates Rab1 deactivation by a non-canonical hydrolytic mechanism. J Biol Chem, 288, 24000–11.

Navaza, J. 1994. AMoRe: an automated package for molecular replacement. Acta Crystallographica Section A Foundations of Crystallography, 50, 157–163.

Petoukhov, M. V., Franke, D., Shkumatov, A. V., Tria, G., Kikhney, A. G., Gajda, M., Gorba, C., Mertens, H. D., Konarev, P. V. & Svergun, D. I. 2012. New developments in the ATSAS program package for small-angle scattering data analysis. J Appl Crystallogr, 45, 342–350.

Romantsov, T., Guan, Z. & Wood, J. M. 2009. Cardiolipin and the osmotic stress responses of bacteria. Biochim Biophys Acta, 1788, 2092–100.

Schmid, E. M., Bakalar, M. H., Choudhuri, K., Weichsel, J., Ann, H., Geissler, P. L., Dustin, M. L. & Fletcher, D. A. 2016. Size-dependent protein segregation at membrane interfaces. Nat Phys, 12, 704–711.

Schneidman-Duhovny, D., Hammel, M., Tainer, J. A. & Sali, A. 2016. FoXS, FoXSDock and MultiFoXS: Single-state and multi-state structural modeling of proteins and their complexes based on SAXS profiles. Nucleic Acids Res, 44, W424–9.

Tesmer, J. J., Berman, D. M., Gilman, A. G. & Sprang, S. R. 1997. Structure of RGS4 bound to AlF4--activated G(i alpha1): stabilization of the transition state for GTP hydrolysis. Cell, 89, 251–61.

Tickle, I. J., Bricogne, G., Flensburg, C., Keller, P., Paciorek, W., Sharff, A. & Vonrhein, C. 2016. Staraniso. Global Phasing Ltd., Cambridge, United Kingdom: http://staraniso.globalphasing.org/cgi-bin/staraniso.cgi.

Treuner-LANGE, A., Macia, E., Guzzo, M., Hot, E., Faure, L. M., Jakobczak, B., Espinosa, L., Alcor, D., Ducret, A., Keilberg, D., Castaing, J. P., Lacas GERVAIS, S., Franco, M., SØGAARD-ANDERSEN, L. & Mignot, T. 2015. The small G-protein MglA connects to the MreB actin cytoskeleton at bacterial focal adhesions. The Journal of cell biology, 210, 243–56.

Treuner-LANGE, A. & Sogaard-ANDERSEN, L. 2014. Regulation of cell polarity in bacteria. J Cell Biol, 206, 7–17.

Vonrhein, C., Flensburg, C., Keller, P., Sharff, A., Smart, O., Paciorek, W., Womack, T. & Bricogne, G. 2011. Data processing and analysis with the autoPROC toolbox. Acta Crystallographica Section D: Biological Crystallography, 67, 293–302.

Wittinghofer, A. & Vetter, I. R. 2011. Structure-function relationships of the G domain, a canonical switch motif. Annu Rev Biochem, 80, 943–71.

Wuichet, K. & Sogaard-ANDERSEN, L. 2014. Evolution and diversity of the Ras superfamily of small GTPases in prokaryotes. Genome Biol Evol, 7, 57–70.

Zeno, W. F., Baul, U., Snead, W. T., Degroot, A. C. M., Wang, L., Lafer, E. M., Thirumalai, D. & Stachowiak, J. C. 2018. Synergy between intrinsically disordered domains and structured proteins amplifies membrane curvature sensing. Nat Commun, 9, 4152.

Zhang, B., Zhang, Y., Wang, Z. & Zheng, Y. 2000. The role of Mg2+ cofactor in the guanine nucleotide exchange and GTP hydrolysis reactions of Rho family GTP-binding proteins. J Biol Chem, 275, 25299–307.

Zhang, Y., Franco, M., Ducret, A. & Mignot, T. 2010. A bacterial ras-like small GTP-binding protein and its cognate GAP establish a dynamic spatial polarity axis to control directed motility. PLoS Biology, 8.

Zhang, Y., Guzzo, M., Ducret, A., Li, Y.-Z. & Mignot, T. 2012. A Dynamic Response Regulator Protein Modulates G-Protein–Dependent Polarity in the Bacterium Myxococcus xanthus. PLoS Genetics, 8, e1002872–e1002872.

